# Nutrient parsimony shapes diversity and functionality in hyper-oligotrophic Antarctic soils

**DOI:** 10.1101/2020.02.15.950717

**Authors:** Marc W. Van Goethem, Surendra Vikram, David W. Hopkins, Grant Hall, Stephan Woodborne, Thomas J. Aspray, Ian D. Hogg, Don A. Cowan, Thulani P. Makhalanyane

**Affiliations:** Centre for Microbial Ecology and Genomics, Department of Biochemistry, Genetics and Microbiology, University of Pretoria, Pretoria 0028, South Africa; Scotland’s Rural College (SRUC), West Mains Road, Edinburgh, EH9 3JG, United Kingdom; Mammal Research Institute, University of Pretoria, Private Bag X20, Hatfield, 0028, South Africa; iThemba LABS, Private Bag 11, WITS, 2050, South Africa; School of Energy, Geoscience, Infrastructure and Society, Heriot-Watt University, Edinburgh, EH14 4AS, United Kingdom; University of Waikato, New Zealand

**Author notes:** Corresponding author: Dr Thulani P. Makhalanyane. Environmental Genomics and Systems Biology Division, Lawrence Berkeley National Laboratory, 1 Cyclotron Rd, Berkeley, CA, 94720, USA.

**Keywords:** Antarctica, carbon cycling, nitrogen cycling, soil stoichiometry, soil respiration

## Abstract

The balance of nutrients in soil is critical for microbial growth and function, and stoichiometric values below the Redfield ratio for C:N:P can negatively affect microbial ecosystem services. However, few studies have assessed the relationships between nutrient balance and biological productivity in extremely nutrient-poor habitats. The Mackay Glacier region of Eastern Antarctica is a hyper-oligotrophic ice-free desert and is an appropriate landscape to evaluate the effects of nutrient deficiency and imbalance on microbial community ecology. In a survey of multiple, widely dispersed soil samples from this region, we detected only low rates of microbial respiration, and observed that C:N:P ratios were well below those required for optimal activity. *In silico* metagenomic and soil isotopic ratio (*δ*^15^N) analyses indicated that the capacity for nitrogen fixation was low, but that soil microbial communities were enriched for soil nitrate assimilation processes, mostly associated with heterotrophic taxa. *δ*^13^C isotope ratio data suggested that carbon dioxide was fixed principally via the Calvin cycle. Genes involved in this pathway were common to all metagenomes and were primarily attributed to members of the dominant soil bacterial phyla: *Bacteroidetes* and *Acidobacteria*. The identification of multiple genes encoding non-photoautotrophic RUBISCO and carbon dioxide dehydrogenase enzymes in both the metagenomic sequences and assembled MAGs is suggestive of a trace-gas scavenging physiology in members of these soil communities.

## Introduction

Soil microorganisms mediate key biogeochemical cycles through an array of complex synergistic interactions [1]. Microbial communities mediate the conversion of key soil elements, including carbon (C) [2], nitrogen (N) [3] and phosphorus (P). Together, these nutrients provide a framework for exploring the balance of organic matter and nutrient availability in the environment. Comparisons with the ideal Redfield atomic ratio of 186:13:1 (C:N:P) for soil, or 60:7:1 for soil biomass [4] have been used to infer resource availability and explain observed variations in soil respiration [5]. In ecosystems close to the cold and arid limits of life, an imbalance in organic matter availability; i.e., values below the Redfield ratio, limit the capacity for microorganisms to mediate biogeochemical cycling [6]. In Antarctic soils, the extremely low levels of C and N, consistent with definitions for very low Redfield ratio values substantially constrain microbial growth and activity [7]. Nevertheless, the limited abundance of higher eukaryotes in continental Antarctica soil habitats [8, 9] ensures that the microbial communities are the dominant suppliers of ecosystem services [10]. The extent to which very low Redfield ratios may limit productivity in these systems has not been explored.

Understanding the functional roles and responses of microbial communities in the warming cryosphere is of considerable contemporary interest. Most studies on continental (non-maritime) Antarctic microbial systems have focused on soils from the McMurdo Dry Valleys in the Ross Dependency [11]. McMurdo Dry Valley soils are typically dominated by oligotrophic bacterial guilds (mainly *Proteobacteria* and *Actinobacteria*) with low representations of *Archaea* and *Eukaryotes* [12]. Microbial activity (measured as respiration rates) in continental Antarctic soils is generally very low [13, 14] per unit of soil, but high when expressed on a microbial biomass specific basis (Hopkins et al. submitted). Microbial respiration significantly influenced by changes in temperature, water bioavailability and substrate availability [15, 16]. Surprisingly, relatively little is known of the composition, diversity and functional traits of microbiomes in other arid Antarctic non-maritime environments such as the Mackay Glacier region [17].

The ∼100km region north of the Mackay Glacier, Eastern Antarctica (Fig. 1) is suggested to be a transitional zone, based on differences in physicochemical features [18] and invertebrate biodiversity compared with surrounding regions [18, 19]. The patchy distribution of micro-invertebrates in the Mackay Glacier region differs substantially from that in the Dry Valleys [20], which is consistent with observations of species patchiness across other transitional zones [21]. Accordingly, we predict that differences in microbial species distribution resulting from soil geomorphic heterogeneity may also support substantially different microbiomes compared to those seen in the Dry Valleys. We predicted that the hyper-oligotrophic status of Mackay Glacier soils would be reflected as imbalanced Redfield ratios, and that major deviations from the ideal ratio could explain respiration rates and unique taxonomic characteristics. To test this hypothesis, we assessed the soil microbial diversity and functional potential of 18 soil communities from sites up to 100km north of the Mackay Glacier, complimented with soil physicochemistry, respiration rates and stable isotope ratios (*δ*^13^C and *δ*^15^N).

**Figure 1.**
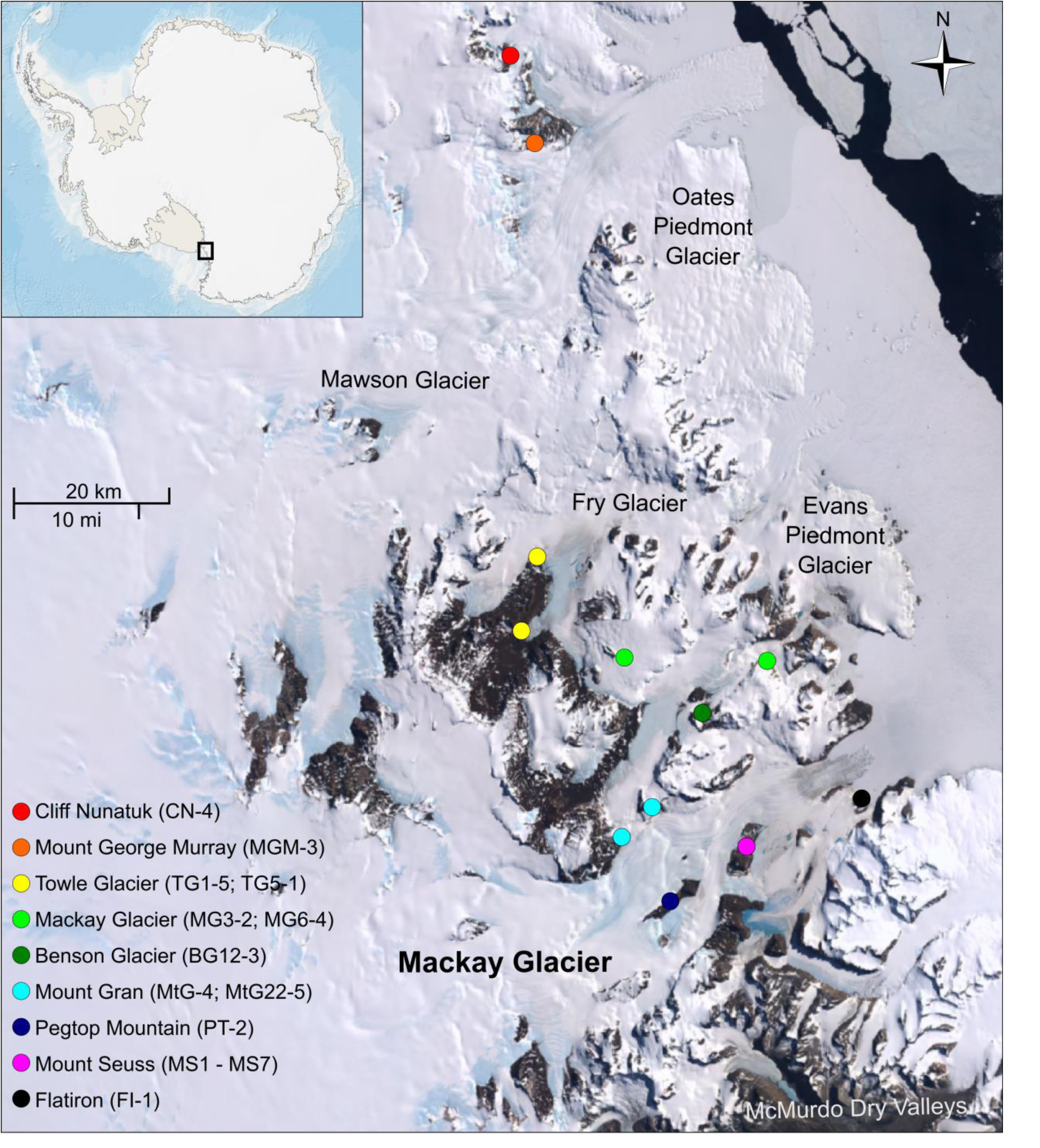
Satellite image of the Mackay Glacier region with the sampling sites indicated. Map of Antarctic is superimposed with the region of interest highlighted. Source: Landsat Image Mosaic of Antarctica (LIMA) Digital Database.

## Materials and methods

### Study site and sampling

The Mackay Glacier (76.52°S 161.45°E) is located to the north of the McMurdo Dry Valleys, Victoria Land, Antarctica. Surface mineral soil samples were recovered from 18 ice-free sites up to ∼100 km north of the McKay Glacier, Eastern Antarctica (Table S1). At each of the 18 sites, approximately 20 g samples were retrieved aseptically from five positions within a 1 m^2^ quadrat (0-5 cm soil depth), providing a total of 90 soil samples. An additional ∼100 g sample was collected at each site for soil physicochemical analysis (*n*=18). Soils were stored in sterile Whirl-Pak bags (Nasco International, Fort Atkinson, WI, USA) and in sterile 50 ml polypropylene Falcon tubes (Grenier, Bio-One) on ice during sampling and transport in the Antarctic, and at - 80°C in the laboratory (Centre for Microbial Ecology and Genomics, University of Pretoria, South Africa) until processed. Sieved soils were analysed for total nitrogen, total carbon and major elements (K^+^, Na^+^, Ca^2+^, and Mg^2+^) using X-ray fluorescence spectrometry on a Philips PW1404 XRF and combustion analysis on a LECO TruSpec® Elemental Determinator. Soil pH was measured using 1:2.5 (mass:volume) soil and deionised water suspensions. Standardised procedures were used for all soil physicochemical analyses at the Stellenbosch Central Analytical Facilities (CAF, Stellenbosch University, RSA).

### Carbon dioxide release measurements

Approximately 10 g (± 0.5) of soil (fresh weight) was incubated at 10°C in the dark for up to 40 days in a conductometric respirometer [22]. Microbial CO_2_ production (respiration) was measured at hourly intervals and output data were used to estimate the soil respiration rates using linear regressions of CO_2_ production over the initial 2-3 days [13].

### Soil Isotopic analyses

For *δ*^13^C analyses, ∼5 g samples were subjected to standard acid pre-treatment to exclude carbonates and then rinsed to neutrality. Samples were then re-dried and ground for carbon isotope ratio analysis while unpretreated samples were used for nitrogen isotope ratio analysis. Soil stable isotope ratios are reported as *δ*^15^N and *δ*^13^C as part per mille (‰) depletion or enrichment of ^15^N or ^13^C, in relation to conventional standards, atmospheric N_2_ and VPDB, respectively. Blanks and standards were analysed at a ratio of 1:12 relative to soil samples, and at the start and end of batch runs. Soil aliquots were analysed in duplicate for *δ*^13^C at the Stable Light Isotope Laboratory (University of Pretoria, South Africa) on a DeltaV isotope mass spectrometer coupled with a Flash EA 1112 series elemental analyser using a Conflo IV (ThermoScientific, USA).

### Nucleic acid extraction and shotgun metagenome sequencing

Duplicate metagenomic DNA extractions were performed on each soil sample from the 18 sites, using an established phenol/chloroform protocol [23]. From each site, pooled samples with the highest DNA concentration and purity (*n*=18) were submitted to a commercial supplier for Illumina-HiSeq sequencing (Mr DNA, Shallowater, TX, USA). Sequencing was performed using paired-ends (2 × 250 bp) for 500 cycles using the HiSeq 2500 Ultra-High-Throughput Sequencing System (Illumina), as per the manufacturer’s instructions.

### Sequence processing and analysis

All sequence data were filtered, trimmed and screened using a combination of in-house scripts and PRINSEQ-lite v0.20.4 [24]. FLASH v1.2.11 was used to merge complimentary forward and reverse reads [25]. We culled low-quality reads and sequences containing ambiguous bases that were not expected to contribute to assembly. The unassembled metagenomes can be found on the MG-RAST server [26] under sample accession numbers 4667018.3 through 4667036. Metagenomic sequences were *de novo* assembled using metaSPAdes v3.9.0 [27] as per a proposed pipeline [28]. Iterative assemblies using increasing *k*-mer lengths generated contigs used in downstream analyses. The quality of each assembled metagenome was evaluated using MetaQUAST v4.3 [29]. Prodigal v2.6.3 was used to predict and extract prokaryotic open reading frames (ORFs) from contigs longer than 200 bp [30]. To provide functional assignments, all ORFs were compared to the NCBI protein non-redundant database at an E value cut-off of 1 × 10^−5^ using DIAMOND v0.7.9.58 [31]. Functional annotations of the ORFs were based on KEGG (Kyoto Encyclopedia of Genes and Genomes) pathways as assigned and visualised in MEGAN5 [32]. Hits corresponding to specific taxa or functions were retained if their bit scores were within 10% of the best bit score. Taxonomic assignments were obtained using the same procedure on all contigs greater than 200 bp in length. All singletons were excluded from analyses.

### Statistical analyses

Bray-Curtis dissimilarities based on Hellinger-transformed relative abundances of microbial genera and functional ORFs identified in each sample were used to calculate dissimilarities between sites. These dissimilarity data were used to create ordinations using non-parametric multi-dimensional scaling (nMDS). We used a redundancy analysis (RDA) within the *vegan* package in R [33] to explore which environmental parameters were significant drivers of microbial community structure within the transition boundary. A permutational non-parametric analysis of variance (PERMANOVA) using the *adonis* function in R was used to test for differences between groups after performing 9999 permutations. Estimates of community richness and diversity were computed using *BiodiversityR* [34], and Nonpareil [35]. Correlations between soil physicochemical variables and community taxonomic data were computed using all Spearman’s rank correlation coefficients (*rho*) that were above 0.6 or below -0.6 and significant (*P* < 0.05).

### Genome binning

Assembled contigs longer than 1.5 kbp were selected for genome binning. Genome binning was performed with the Anvi’o v4 platform [36] and CONCOCT v1.0.0 binning software [37]. Complete bins with the lowest contamination were selected for the next round of refinement using sample-specific read alignment, extraction and reassembly using the SPAdes v3.9.0 assembler [38]. Complete or near-complete bins were selected for refinement in the Anvi’o pipeline. Bin completeness, contamination and strain heterogeneity were assessed at each step using CheckM [39]. The bins with the least contamination and heterogeneity were selected for functional annotation and phylogenetic analysis. The high-quality bins were used to identify single copy marker genes using AMPHORA2 [40]. Out of 31 single copy marker genes, 7 and 11 genes were common in all the reference genomes and target bins belonging to *Chitinophagales* and *Acidobacteria*, respectively. The selected protein sequences were aligned, trimmed and a final concatenated tree was generated using the neighbour-joining method [41] in MEGA7 [42]. Contigs were aligned with the sample-specific reads using bowtie2 aligner [43] to quantify the average coverage in the respective samples. Functional annotation of reconstructed genomic bins was performed with the RAST [44] and KAAS [45] servers. Circular genome visualisation was performed using DNAPlotter [46].

## Results

The location of samplings sites is shown in Fig. 1, at altitudes ranging from 157 – 1,109 m above sea level (Table S1). Physicochemical analyses showed that the soils were slightly alkaline (pH 7.5 – 8.7) and extremely oligotrophic, with soil carbon (0.1 – 0.15%) and nitrogen (0.008 – 0.042%) levels close to the accurate detection limits (Table 1). Soil C:N ratios varied considerably (2.86 – 15), and most were below the ideal Redfield ratio of 6.6:1, suggesting that nitrogen is severely limiting at most sample sites. Surface soil phosphorus (P) concentrations ranged widely across all samples, from 9 – 220 ppm (mean 41.26 ppm). Combined elemental ratios of ∼60:0.05:0.01 were markedly lower than the ideal Redfield ratio of 60:7:1 for optimal soil microbial community activity and biomass [4, 5]. All samples were also characterised by low major cation contents (Table S1).

**Table 1.** Values of *δ*^13^C and *δ*^15^N (‰), soil carbon, nitrogen and phosphorus, and stoichiometric ratios for Mackay Glacier soils.

| Sample ID | $\delta^{13}\text{C}$ (‰) | $\delta^{15}\text{N}$ (‰) | % C | % N | P (ppm) | C:N Ratio | N:P Ratio | C:P Ratio |
| --- | --- | --- | --- | --- | --- | --- | --- | --- |
| BG12-3 | -31.34 | -10.36 | 0.10 | 0.020 | 11 | 5.0 | 0.0018 | 0.009 |
| CN-4 | -26.42 | -9.755 | 0.14 | 0.016 | 12 | 8.75 | 0.0013 | 0.012 |
| FI-1 | -27.38 | -0.406 | 0.12 | 0.042 | 20 | 2.88 | 0.0021 | 0.006 |
| MG3-2 | -28.46 | -8.084 | 0.10 | 0.020 | 9 | 5.0 | 0.0022 | 0.011 |
| MG6-4 | -29.95 | -5.786 | 0.12 | 0.016 | 18 | 7.5 | 0.0009 | 0.007 |
| MGM-3 | -26.50 | -2.314 | 0.12 | 0.032 | 25 | 3.75 | 0.0013 | 0.005 |
| MS1-1 | -28.35 | -1.713 | 0.15 | 0.032 | 22 | 4.69 | 0.0015 | 0.007 |
| MS2-2 | -26.47 | 0.019 | 0.12 | 0.016 | 24 | 7.5 | 0.0007 | 0.005 |
| MS3-5 | -28.14 | -6.605 | 0.10 | 0.017 | 14 | 5.88 | 0.0012 | 0.007 |
| MS4-1 | -26.53 | -5.020 | 0.12 | 0.021 | 17 | 5.71 | 0.0012 | 0.007 |
| MS5-1 | -26.82 | -2.401 | 0.10 | 0.010 | 60 | 10.0 | 0.0002 | 0.002 |
| MS6-5 | -24.84 | 2.720 | 0.12 | 0.017 | 15 | 7.06 | 0.0011 | 0.008 |
| MS7-5 | -28.69 | -8.247 | 0.13 | 0.026 | 23 | 5.0 | 0.0011 | 0.006 |
| MtG-4 | -23.72 | -6.891 | 0.10 | 0.024 | 15 | 4.17 | 0.0016 | 0.007 |
| MtG22-5 | -23.45 | -0.793 | 0.10 | 0.028 | 19 | 3.57 | 0.0015 | 0.005 |
| PT-2 | -27.01 | -11.58 | 0.12 | 0.008 | 4 | 15.0 | 0.0020 | 0.030 |
| TG1-5 | -28.91 | -11.78 | 0.12 | 0.028 | 12 | 4.29 | 0.0023 | 0.010 |
| TG5-1 | -27.29 | -8.670 | 0.12 | 0.019 | 11 | 6.32 | 0.0017 | 0.011 |

In dark-incubated microcosms, CO_2_ production increased linearly over 2-3 days for most samples, giving low but reliable rates of microbial respiration (average value of 0.407 μg C g^-1^ soil d^-1^). CO_2_ flux values from the soils ranged widely from 0.075 to 0.938 μg Cg^-1^ soil d^-1^ (Table S2). Carbon stable isotope ratios (*δ*^13^C range; -23.5 to -31.3‰) were consistent with C metabolism via the Calvin-Benson-Bassham (CBB) cycle (Table 1). Nitrogen stable isotope ratios indicated a predominance of *in situ* soil nitrate assimilation processes [13], with only two samples providing evidence of nitrogen fixation (*δ*^15^N range, 2.7 to -11.8‰).

We used shotgun metagenomic sequencing to evaluate the community structure and functional potential of the soil microbiomes (a summary of sequencing statistics is provided in Table S3). Sequencing produced ∼50 Gbp of high-quality sequence data, sufficient to describe more than 95% of the soil microbial diversity (Fig. S1). After sequence assembly, we found that bacterial contigs dominated all metagenomes (94 – 99% of contigs), while Eukaryotic contigs ranged from 0.2 to 5.0%, and *Archaea* represented less than 1.5% of the community. Contigs assigned to viruses were extremely rare (absent to 0.1%), but those identified were principally assigned as *Caudovirales* [47].

*Bacteroidetes* (34 – 73%), *Acidobacteria* (4 – 26%), *Proteobacteria* (6 – 18%) and *Cyanobacteria* (2 – 10%) were the dominant bacterial phyla in all soil communities (Fig. 2*A*, Table S4). Surprisingly, all communities were dominated by members of the *Bacteroidetes* and *Acidobacteria*, which together accounted for more than 68% of contigs longer than 200 bp. Other common community members included *Actinobacteria, Firmicutes, Verrucomicrobia* and *Chloroflexi.*

**Figure 2.**
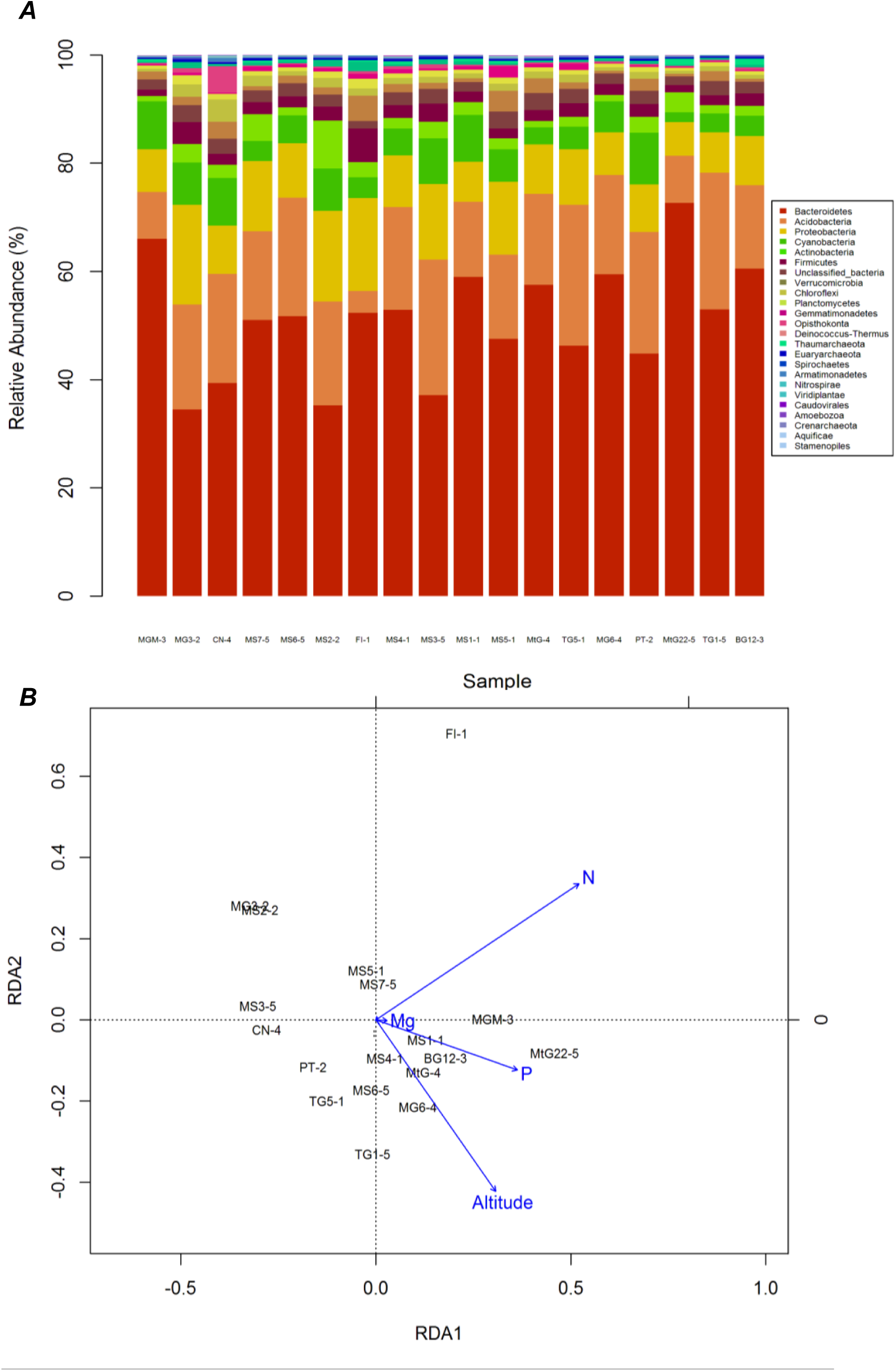
Microbial community composition and functional patterns across the 18 soil communities. (A) Metagenomic taxonomy classified by major phyla. Samples are arranged in order of increasing altitude, from 157 m.a.s to 1109 m.a.s, left to right. (B) Redundancy analysis (RDA) bi-plot of community structure and abiotic parameters (left). Only the abiotic features that significantly explained variation in microbial community structure are fitted onto the ordination (blue arrows). The length of the arrow is proportional to the rate of change. The direction of the arrow corresponds to the direction of maximum change of that variable.

Overall, soil microbial composition was significantly influenced by site altitude (ANOSIM, *P* < 0.005) and soil N content (*P* < 0.005), with more subtle differences attributed to soil P (*P* < 0.02) and magnesium (*P* < 0.05) content (Fig. 2*B*). The mean estimates of inferred species diversity (alpha diversity: *α*) of Mackay Glacier region soils (mean *α* = 452.5, *n* =18) were slightly lower than those from McMurdo Dry Valley soil metagenomes that are publicly available on MG-RAST (mean *α* = 557.6; *n* = 11, *P* > 0.05).

We analysed the functional potential of each Mackay glacier soil community metagenome to gain a deeper understanding of the capacity for microbial biochemical cycling at challenging C:N:P ratios (Table S5). We found that 17 of the 18 metagenomes harboured signature genes for CO_2_ fixation via the Calvin-Benson-Bassham (CBB) cycle, which was consistent with our *δ* ^13^C data trends. Just over half of the ribulose-1,5-bisphosphate carboxylase oxygenase (RuBisCO) (*n*=79) and phosphoribulokinase (*prk*B) (*n*=48) genes identified were assigned to *Cyanobacteria* (Fig. 3), which are the dominant photosynthetic carbon fixers in most desert soils [10, 48, 49], and were relatively common here. The remaining 83 gene assignments belonged to potentially chemosynthetic guilds, including *Actinobacteria, Bacteroidetes, Proteobacteria* and *Planctomycetes* (Table S6), which may together serve as important primary producers in soil, and is consistent with the known importance of alternative primary producers in Antarctic soils [50]. Genes involved in the Arnon-Buchanan cycle were also common (*kor*AB, 2-oxoglutarate: ferredoxin oxidoreductase; *n*=1652), suggesting an additional active mechanism of CO_2_ fixation. Carbon monoxide oxidation genes (*cox*LMS) mainly belonged to *Bacteroidetes* (*n*=218) and *Acidobacteria* (*n*=62), which potentially offers a viable alternative carbon scavenging mechanism to carbon fixation in this carbon limited environment.

**Figure 3.**
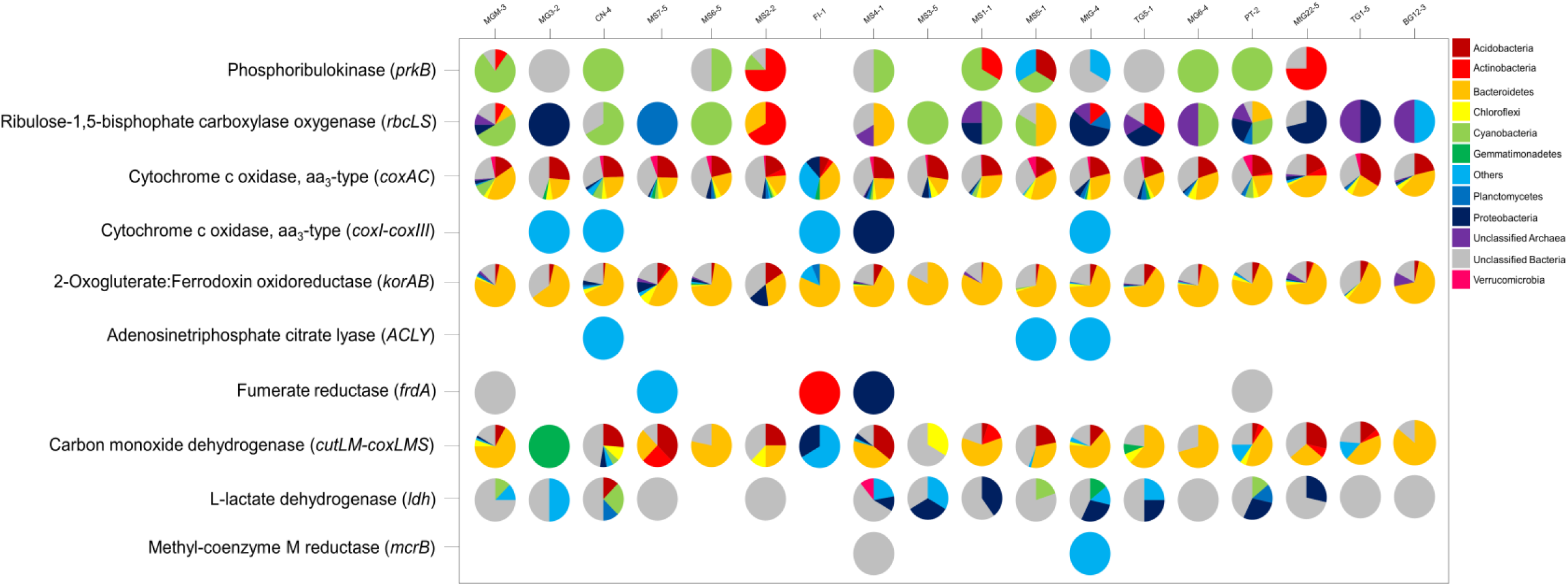
Taxonomic assignments of functionally-annotated ORFs for genes involved in carbon cycling classified at the phylum level for the 18 metagenomes. Samples are arranged by increasing altitude.

The depleted *δ* ^15^N ratios (2.7 to -11.8‰) suggest that nitrogen fixation was not a dominant N-assimilation pathway in the Mackay Glacier region soil microbiomes [51]. This was consistent with the observation that indicator genes for N_2_ fixation (*nif*DHK) were uncommon (6 significant matches across all metagenomes), despite the presence of phylogenetic signals for heterocystous *Cyanobacteria* in all soil samples. Instead, the assembled metagenomes were rich in genes from nitrate assimilation and nitrogen mineralization pathways, most of which were assigned to the *Bacteroidetes* and *Acidobacteria* (Table S6).

Indicator genes for respiratory ammonification (*nrf*A) were most common in samples collected at higher altitudes (>550 m above sea level). In contrast, *nar*GH genes that are indicative of nitrate reduction were found only in low altitude (coastal) soil communities (Fig. 4). The subsequent step in the nitrogen cycle, denitrification of NO_3_^-^ to N_2_ gas is indicated by *nor*BC genes, which were assigned to *Bacteroidetes* and *Verrucomicrobia*, although these were not present in all metagenomes. Ammonia oxidation pathway genes, *amo*ABC, were taxonomically assigned to known ammonia-oxidising guilds within the *Thaumarchaeota* and *Proteobacteria*.

**Figure 4.**
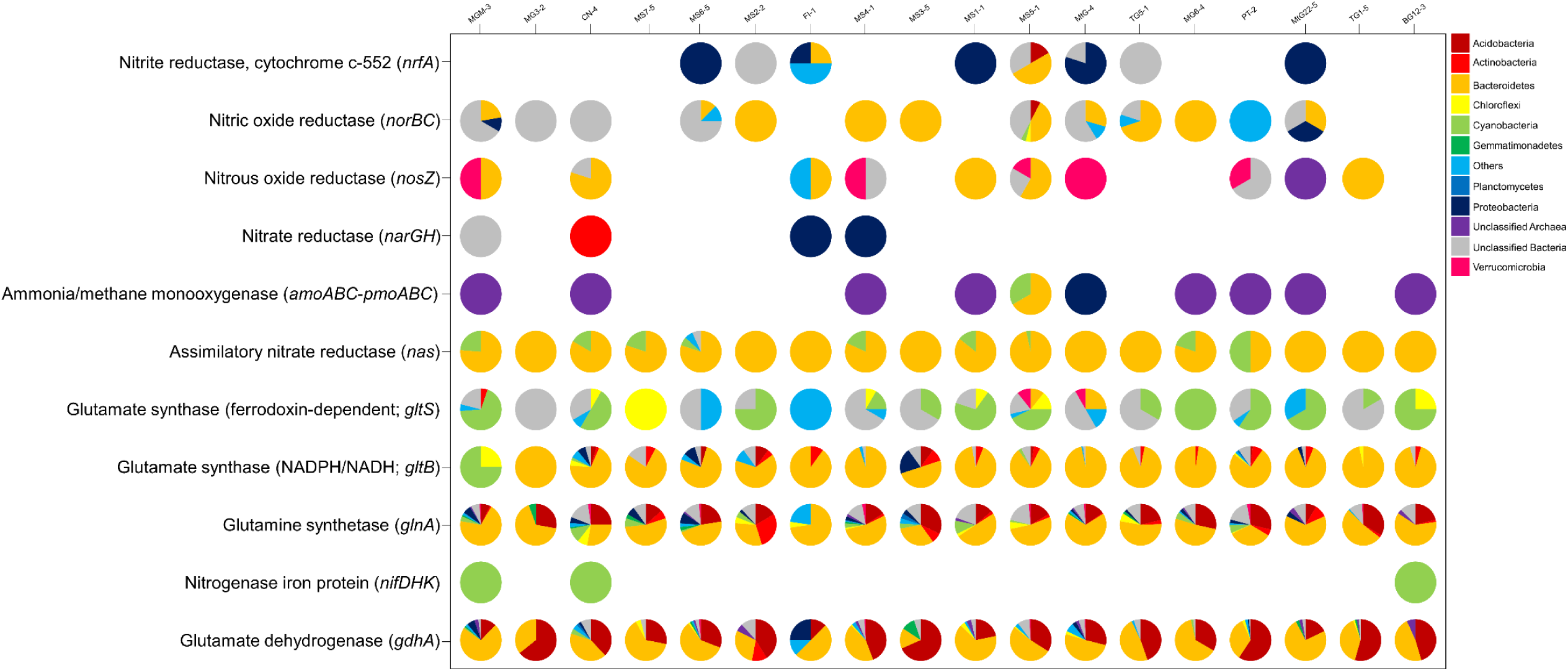
Taxonomic assignments of functionally-annotated ORFs for genes involved in nitrogen cycling classified at the phylum level for the 18 metagenomes. Samples are arranged by increasing altitude.

We reconstructed and refined seven high-quality draft genomes of dominant taxa from an initial set of 80 metagenome-assembled genomes (MAGs). Here we describe the functional attributes of the two most complete draft genomes, belonging to the members of the dominant phyla *Bacteroidetes* and *Acidobacteria* (Table S7). Both reconstructed genomes constituted near-complete MAGs; i.e., ≥ 90% completeness and ≤ 5% contamination [39, 52].

The first MAG, designated as *Chitinophagales* bacterium 62-2, comprised 4.9 Mbp of sequence in 756 contigs (92.5% completeness). This MAG was phylogenetically similar to members of the genus *Segetibacter* (family *Chitinophagaceae*; *Bacteroidetes*; Fig. 5*B*). Contigs belonging to the genus *Segetibacter* were abundant in all metagenomes, with a range of 0.5 – 3.9%, and a mean relative abundance of 2.1%. The *Chitinophagales* bacterium 62-2 MAG was smaller than the closest sequenced isolate, *Segetibacter koreensis* (6.1 Mbp), which was obtained from temperate soil [53]. Nonetheless, the MAG retained many of the genomic attributes for an aerobic chemosynthetic lifestyle, including genes for substrate transport, glycan import and pathways for glucose fermentation and oxidation, as well as photo-receptor genes (Fig. 5*A*).

**Figure 5.**
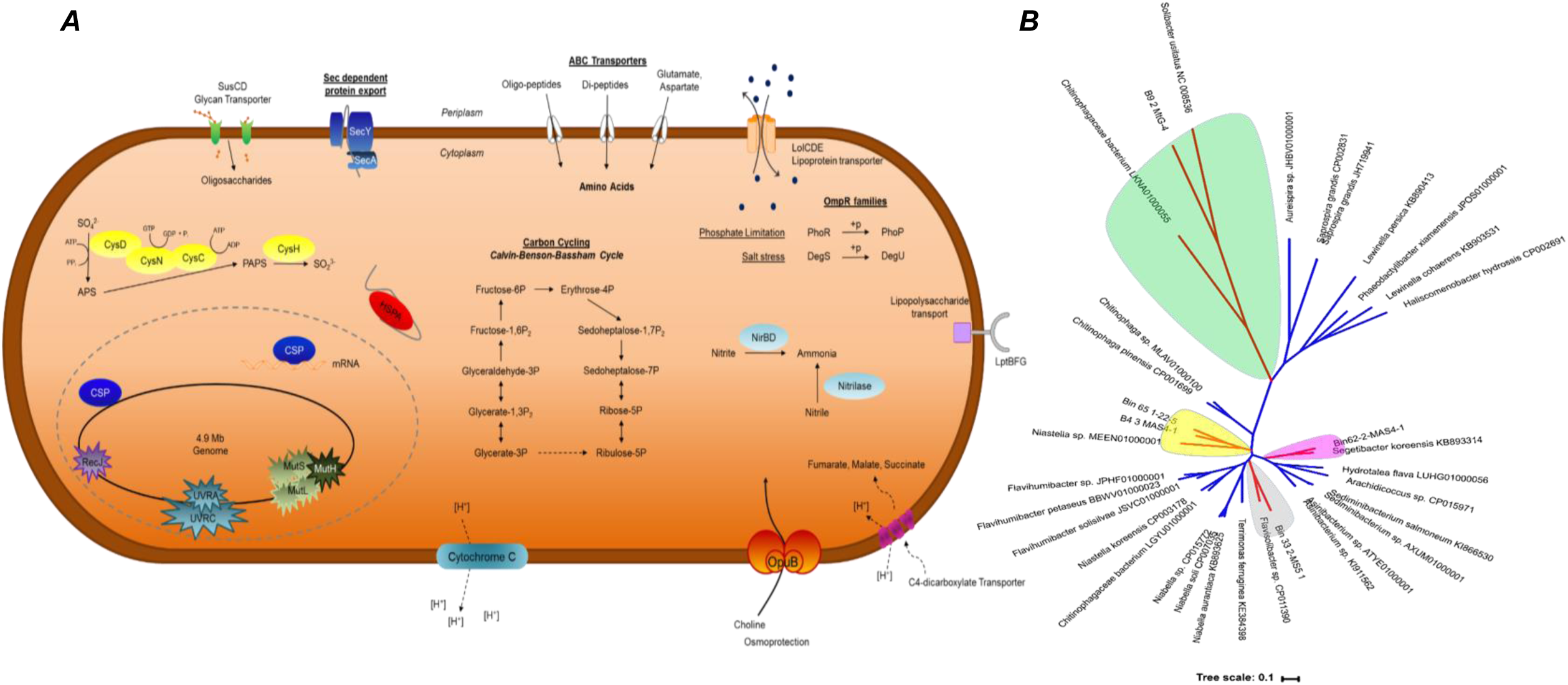
(*A*) Schematic overview of the *Bacteroidetes* (Bin_62-2_MS4-1) MAG indicating important genes involved in C and N cycling. Furthermore, various stress response adaptations, including those involved in DNA repair and osmoprotection are indicated as well as transmembrane transporters, mechanisms of sulphur cycling and genes associated with cold stress responses. The assignments were based the RAST and KAAS web servers. (*B*) Phylogenetic tree placing Bin_62-2_MS4-1 within the *Segetibacter* (Chitinophagales).

The second MAG, *Acidobacteria* bacterium 28-9, showed a high level of phylogenetic relatedness to the genus *Pyrinomonas* (*Acidobacteria*), and comprised ∼3.7 Mbp of sequence in 119 contigs with 92.3% completeness (Fig. 6*A, B*). Members of the genus *Pyrinomonas* contributed an average of 5.4% of contigs in all metagenomes (range 0.5–11.9%). The *Acidobacteria* bacterium 28-9 genome had a low G+C content (45.7%), compared to both polar and temperate *Acidobacteria* isolates; *Granulicella tundricola, Acidobacterium capsulatum* and *Ellin345* (∼60.3%) [54, 55]. In comparison to temperate *Acidobacteria* genomes, we found very few genes for cellular motility and chemotaxis, and no genes encoding flagellar proteins, despite their ubiquity in soil isolates of this taxon.

**Figure 6.**
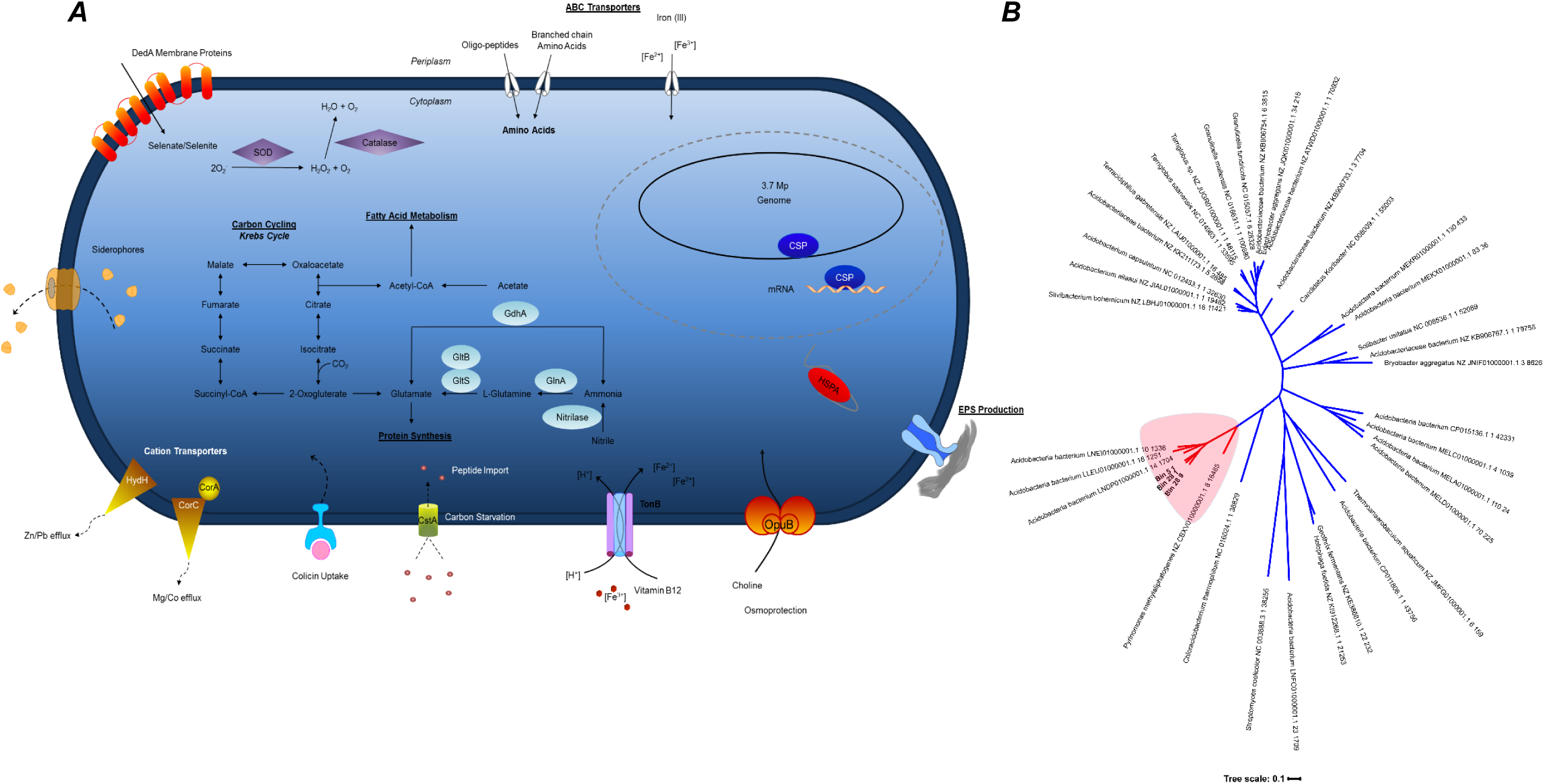
(*A*) Schematic overview of the *Acidobacterium* (Bin_28-9_MtG-4) MAG indicating important genes involved in C and N cycling. Furthermore, various stress response adaptations, including those involved in DNA repair, osmoprotection and heavy metal efflux and genes associated with cold stress responses. The assignments were based the RAST and KAAS web servers. (*B*) Phylogenetic tree placing Bin_28-9_MtG-4 within the *Pyrinomonas* (Acidobacteria).

The genome of *Acidobacteria* bacterium 28-9 encoded an aerobic, heterotrophic metabolism including a near-complete Krebs cycle (Fig. 6*A*), genes for substrate degradation (including amino acids, sugars and complex organic polymers). We also identified a range of psychrotolerance and stress response mechanisms encoded in the *Acidobacteria* bacterium 28-9 genome. These include genes for exopolysaccharide (EPS) production, cold-shock proteins (CspA), superoxide dismutase (SOD) and catalase.

Pathways for environmental amino acid acquisition were common to both MAGs, and probably represent energy conservation strategies that reduce the high metabolic cost of *de novo* amino acid biosynthesis. Both MAGs also encoded heavy metal resistance proteins for cobalt, zinc and mercury export. All genes for the complete trehalose biosynthesis pathway, a well-known desiccation resistance mechanism, were present in both draft genomes.

## Discussion

Desert soils typically have widely varying chemical compositions and ratios of C:N:P [56, 57]. All soil samples in this study were characterised by extremely low C and N contents but high soil P. It is possible that the very low soil C-substrate availability and the extreme microenvironmental characteristics within these soils could have limited the spectrum of *Cyanobacteria* with the capacity to fix atmospheric N via diazotrophy [58, 59]. Here, we observed microbial communities that exist below the C:N:P Redfield ratio for balanced microbial growth (60:7:1). The extreme physiological constraints imparted by the C:N:P stoichiometric imbalance (∼6200:9:1) almost certainly limit biogeochemical cycling within this region of the Antarctic continent. Other continental Antarctic soils, i.e. those studied in the McMurdo Dry Valley region, have biochemical balances much nearer the predicted ratio for optimal microbial growth (10:1:1), although these ratios vary widely according to soil type, soil age, proximity to lacustrine deposits and the exchange of soil nutrients facilitated by liquid water [6].

The rates of soil microbial respiration in the Mackay Glacier region were up to an order of magnitude lower than some Dry Valley soils (0.9 – 4 μg C/g^-1^ soil/d^-1^) [14, 60]. Heterotrophic respiration rates are sensitive to both moisture content and temperature [16, 61], and our data show that limited substrate availability is a substantial constraint on microbial respiration regardless of site altitude [15, 60]. The C and N concentrations are very low compared to some Dry Valley regions known for their oligotrophic status, including the low altitude Miers [62] and Wright Valleys [63], and the high altitude Beacon Valley [64]. We infer that the very low respiration rates but high inferred phylogenetic diversity in these soils is a likely consequence of capturing a high proportion of dormant cells in our analysis. Dormancy is a common strategy for microbial survival in hyperarid soils, whereby microbes suspend their metabolism until conditions improve [65]. In Antarctica, cellular metabolism is undoubtedly low compared to warmer climates, yet our data provide evidence for ongoing nutrient cycling processes.

Our carbon isotopic data were consistent with values documented from previous analyses of Dry Valley soils [13]. The soil *δ* ^13^C values were in line with the conclusion, from our metagenomic sequence data, that the Calvin cycle is probably the dominant carbon fixation mechanism in oligotrophic Antarctic soils [66]. The *δ*^15^N values provided little evidence of *in situ* nitrogen fixation, in contrast to other dryland soil environments [67, 68]. This may be the result of community composition or unbalanced soil nutrient stoichiometry, in particular the limitation of substrate C that is essential for N fixation [6, 59]. Moreover, the *δ* ^15^N data suggested a prevalence of nitrate assimilation, which is catalysed by nitrate reductase enzymes (Table 1; Fig. S4). Soil nitrate is depleted of ^15^N in Antarctic soils [13], and our data support previous findings that soil nitrate in the desert sub-surface may be an essential reservoir of bioavailable nitrogen used by microorganisms to overcome nitrogen deficiency [69]. Interestingly, a recent metatranscriptomic study of microbial functionality in desiccated hot desert soils [70] showed high levels of nitrate reductase gene expression but little or no nitrogenase gene expression, suggesting that nitrate was also the primary source of metabolic N in this edaphic environment.

High NO_3_^-^ assimilation rates in arid soils are also associated with very low rates of biotic denitrification [69]. In our metagenomic surveys, genes for denitrification (*nor*BC, *nos*Z) were almost completely absent from all communities, suggesting that only a minor fraction of soil N is lost from the system via microbial-mediated nitrate reduction. Other contributions to the nitrogen cycle, by ammonification and ammonia oxidation should not be ignored, and recent findings have hinted at their importance in depauperate soil systems [71]. Combined these lines of evidence suggest that Mackay Glacier soils are major sinks for nitrate yet minimal sites for denitrification. However this hints that NO_3_^-^ flux out of these soils may occur under different environmental cues including water receipt or temperature when denitrification, and possibly nitrogen fixation, may become predominant nitrogen pathways [66, 72].

Our redundancy analysis indicated that site altitude and soil nitrogen content were the most important drivers of microbial taxonomic diversity patterns within these soils. Soil N and altitude also influenced viral guilds in Mackay Glacier region soils [47], and it is likely that the biotic relationship between viruses and their cognate hosts also shapes microbial community structure in these environments, as indicated in thawing permafrost communities from the Arctic [73, 74]. Abiotic features such as soil pH, fertility and moisture content, are known to explain compositional differences between soil communities in the Antarctic [62, 75, 76], and elsewhere [77]. Differences in water availability correlate strongly with site altitude [78], which possibly explains the significant negative correlation between altitude and cyanobacterial relative abundance observed here (Fig. S2*A*). *Cyanobacteria* are more sensitive to water availability than many other taxa [79-81]. In continental Antarctica, temperatures decrease with increasing altitude, which reduces the active zone (seasonal melting permafrost) to a point where the ice-to-liquid water-to-gaseous water transition is reduced to a solid-to-gas transition. At this point, bioavailable water is insufficient to support soil microbial communities [82].

Our extensive metagenomic characterisation of the microbial communities indicated that the Mackay Glacier region soils were fundamentally distinct from those of the more southern McMurdo Dry Valleys. In the former, most contigs and 16S rRNA genes were affiliated with *Bacteroidetes* and *Acidobacteria*. In contrast, *Actinobacteria* and *Proteobacteria* are typically dominant bacterial phyla in the Dry Valleys, with *Bacteroidetes* and *Acidobacteria* comprising a relatively minor fraction of sequences [64, 83-85]. For example, a recent comprehensive NGS analysis of soil communities from Victoria Valley showed a dominance of *Actinobacteria* and *Gemmatimonadetes*, with only minor representation from *Acidobacteria* and *Bacteroidetes* [86].

The ecological significance of these differences in the dominant taxonomic groups is unclear, but the identification of certain chemosynthetic genes, particularly those encoding non-photoautotrophic RuBisCO and carbon monoxide dehydrogenase, with affiliations to the phylum *Acidobacteria*, may provide a valid clue. Several recent studies [50, 87, 88] have suggested that trace gas (H_2_, CO) scavenging is an important energy acquisition physiology in extremely oligotrophic soils, and that members of both the *Bacteroidetes* and *Acidobacteria* are implicated in this process [87].

The assembly of representative MAGs from the two dominant phyla, *Bacteroidetes* and *Acidobacteria*, supported this view. The implied physiology of both organisms indicated a specialised mixed chemosynthetic and heterotrophic lifestyle, with a range of adaptations to low substrate concentrations. Consequently, we found that both MAGs encoded genes for oxygen-dependent [NiFe]-hydrogenase metallocentre assembly proteins (*hyp*) and CO dehydrogenase maturation factors (*cox*F) indicating potential alternative chemosynthetic pathways [88]. The gene analysis also suggested that both organisms have the capacity to import amino acids, peptides and a range of carbohydrate substrates. *Bacteroidetes* are typically specialist degraders of high molecular weight compounds [89] and *Acidobacteria* are metabolisers of complex organic substrates [90].

*Acidobacteria* are common soil colonists of oligotrophic habitats [55], and their dominance in these low carbon soils is consistent with *K*-strategist ecology, in which slow-growth is favoured over rapid replication rates [91]. Genome reduction has also been observed in bacteria living in nutrient-limited environments (Baumgartner *et al* 2017). The reduced genome size of Chitinophagales bacterium 62-2 MAG, compared to its closest homologue, is suggestive of such an evolutionary effect in the Antarctic hyper-arid and oligotrophic soils.

Determinants of psychrotolerance and stress mitigation strategies were also widespread in both assembled genomes. Genes involved in cryoprotection included those encoding the production of cryoprotective agents such as betaine and trehalose, which act by lowering the freezing point of the cytoplasm [92]. Genes for exopolysaccharide production were present in both MAGs. *De novo* EPS production is triggered by low temperatures [93] and desiccation [94], and is a fundamental microbial survival strategy in many extreme environments, where EPS structured biofilms provide protection against desiccation and UV irradiance, enhance conductivity and support microbial growth [95].

The repertoire of functional genes detected, the measurements of microbial activity and the isotope data are all indicative of functional microbial communities even under the harsh environment conditions, consistent with many previous studies. The fact that these complementary and mutually confirmatory signals have been detected under conditions that would be considered the be stoichiometrically disadvantages suggests two, non-mutually exclusive, possibilities. First, the soil microbial communities could be especially well adapted to the extreme nutrient conditions, perhaps by very efficient recycling and nutritional parsimony. Second, the chemical speciation of the nutrient elements and physical interactions between nutrient elements and the soil physical components in these Antarctic soils differ from many other soils so that larger fractions of the soil N and P pools are available to the microorganisms.

## Acknowledgements

This research was supported by the South African National Research Foundation (Grant ID. 97891 MWVG, 99320 TPM) and the South African National Antarctic Program (SANAP Grant ID 93074). SV received support from the Claude Leon Foundation. We also acknowledge the University of Pretoria (Genomics Research Institute and Research Development Program). Field logistic and financial support were provided to IH and DAC by Antarctica New Zealand and the New Zealand Antarctic Research Institute (NZARI).

## Author contributions

T.P.M., D.A.C., and I.D.H. designed the research; M.W.V.G., G.H., T.J.A. performed the analyses; M.W.V.G., S.V., D.W.H., and S.W. analysed the data; M.W.V.G., S.V., D.W.H., S.W., I.D.H., D.A.C. and T.P.M. wrote the paper.

## Data availability

The unassembled metagenomes are available on the MG-RAST server under sample accession numbers 4667018.3 - 4667036. The contigs have been deposited in the NCBI database and are available under the BioProject ID: PRJNA376086.

