## Supplementary Materials for "Nutrient parsimony shapes diversity and functionality in hyper-oligotrophic Antarctic soils"

**Running title:** Effects of nutrient stoichiometry in Antarctic soils

**Authors:** Marc W. Van Goethem^1§^, Surendra Vikram^1^, David W. Hopkins^2^, Grant Hall^3^, Stephan Woodborne^3,4^, Thomas J. Aspray^5^, Ian D. Hogg^6^, Don A. Cowan^1^ and Thulani P. Makhalanyane^1*^

**Affiliations:**

^1.^Centre for Microbial Ecology and Genomics, Department of Biochemistry, Genetics and Microbiology, University of Pretoria, Pretoria 0028, South Africa

^2.^ Scotland’s Rural College (SRUC), West Mains Road, Edinburgh, EH9 3JG, United Kingdom

^3.^ Mammal Research Institute, University of Pretoria, Private Bag X20, Hatfield, 0028, South Africa ^4.^ iThemba LABS, Private Bag 11, WITS, 2050, South Africa

^5.^ School of Energy, Geoscience, Infrastructure and Society, Heriot-Watt University, Edinburgh, EH14 4AS, United Kingdom

^6.^ University of Waikato, New Zealand

^§^ Present address: Environmental Genomics and Systems Biology Division, Lawrence Berkeley National Laboratory, 1 Cyclotron Rd, Berkeley, CA, 94720, USA

***Corresponding author:**	Dr Thulani P. Makhalanyane

**Conflict of Interest:** The authors declare they have no conflict of interest.

**Funding Sources:** This work was supported by the South African National Antarctic Program funding instrument of the National Research Foundation.

**Subject Category:** Microbial ecology and functional diversity of natural habitats **Supplementary Figure 1.** Rarefaction curves indicating the proportion of microbial diversity sequenced in our study based on read redundancy and sequencing effort. Nonpareil estimates sequencing coverage based on forward read redundancy of filtered (high-quality) sequences.

**Supplementary Figure 2.** Correlation plots indicating significant (*P* < 0.05) positive (blue) and negative (red) interactions between soil physicochemical features and (*A*) microbial phyla, or (*B*) KEGG-assigned subsystem processes.

**Supplementary Figure 3.** (*A*) Draft *Segetibacter* genome and (*B*) draft *Pyrinomonas* genome. Features correspond to concentric circles, beginning with the outermost circle. (1) DNA (contigs). (2) Genes on the forward strand. (3) Genes on the reverse strand. (4) tRNAs. (5) Genes encoding hypothetical proteins and rRNAs. (6) GC-bias. (7) GC-skew.

**Supplementary Figure 4.** Isotope bi-plot for organic materials of Mackay Glacier soils. The values plotted are mean *δ*^13^C and *δ*^15^N values.

**Supplementary Table 1.** The soil physicochemical and environmental characteristics of the 18 sites along the Mackay Glacier ecotone.

**Supplementary Table 2.** Respiration rates (soil CO_2_ release).

**Supplementary Table 3.** Soil metagenome sequencing statistics.

**Supplementary Table 4.** The relative abundance of major phyla across the 18 soils in arranged in order of increasing altitude. The numbers in cells indicate percentage abundance. The cell colours are based on relative abundances with the highest values in red.

**Supplementary Table 5.** Functional genes analysed for carbon and nitrogen cycling.

**Supplementary Table 6.** Taxonomic assignments of functional genes involved in C and N turnover from each metagenome.

**Supplementary Table 7.** Characteristics of the 7 draft genomes reconstructed in this study.

**Supplementary Figure 1.** Rarefaction curves indicating the proportion of microbial diversity sequenced in our study based on read redundancy and sequencing effort.

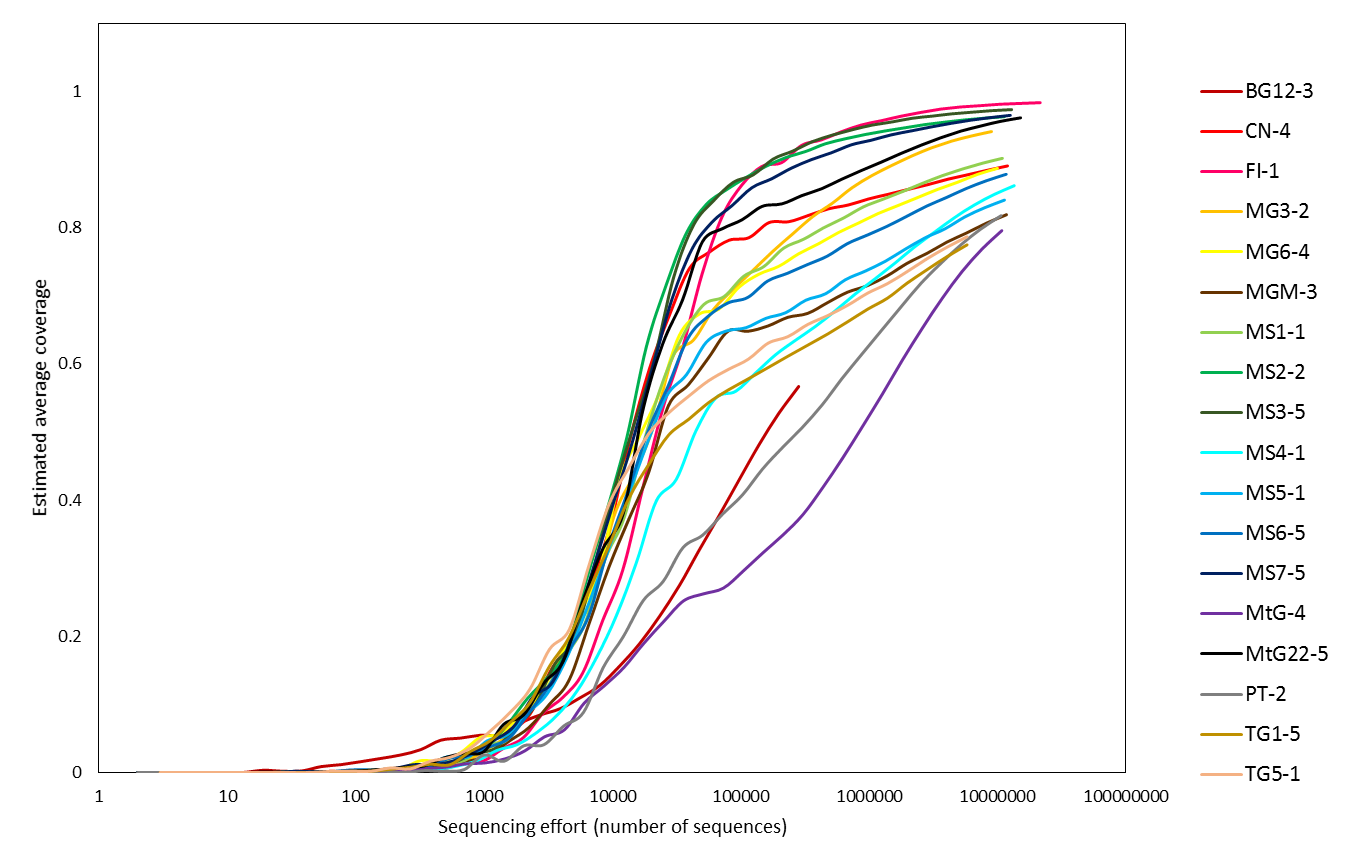

**Supplementary Figure 2.** Correlation plots indicating significant (*P* < 0.05) positive (blue) and negative (red) interactions between soil physicochemical features and (*A*) microbial phyla, or (*B*) KEGG-assigned subsystem processes.

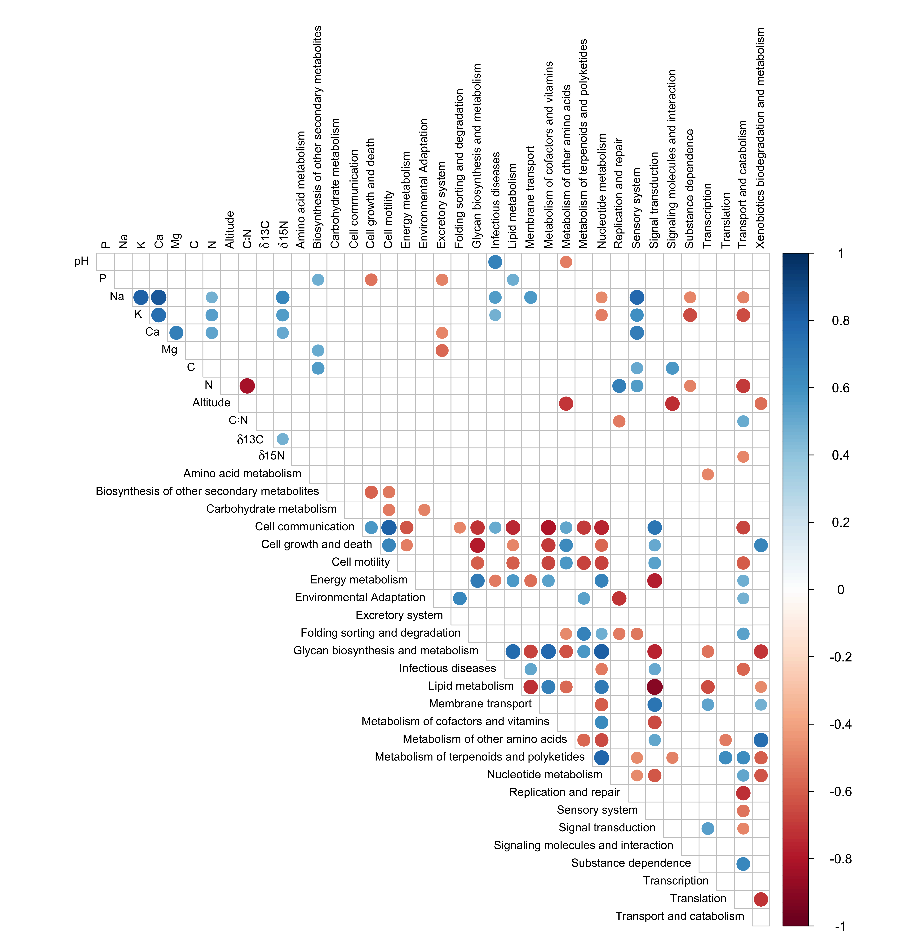

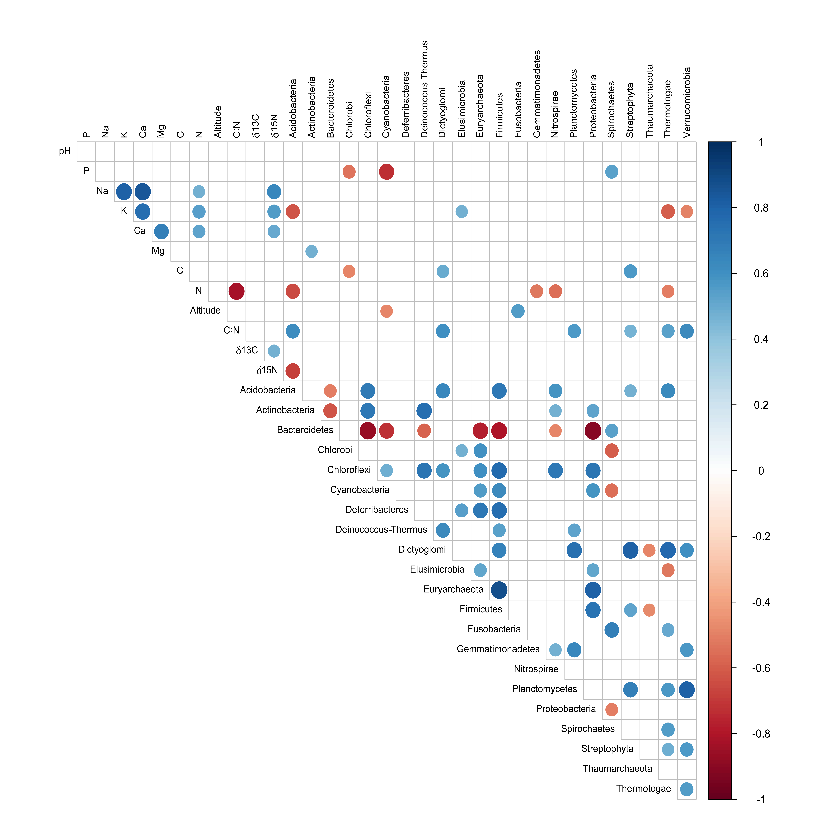

**B**

**A**

**Supplementary Figure 3.** (*A*) Draft *Segetibacter* genome and (*B*) draft *Pyrinomonas* genome. Features correspond to concentric circles, beginning with the outermost circle. (1) DNA (contigs). (2) Genes on the forward strand. (3) Genes on the reverse strand. (4) tRNAs. (5) Genes encoding hypothetical proteins and rRNAs. (6) GC-bias. (7) GC-skew.

**A**

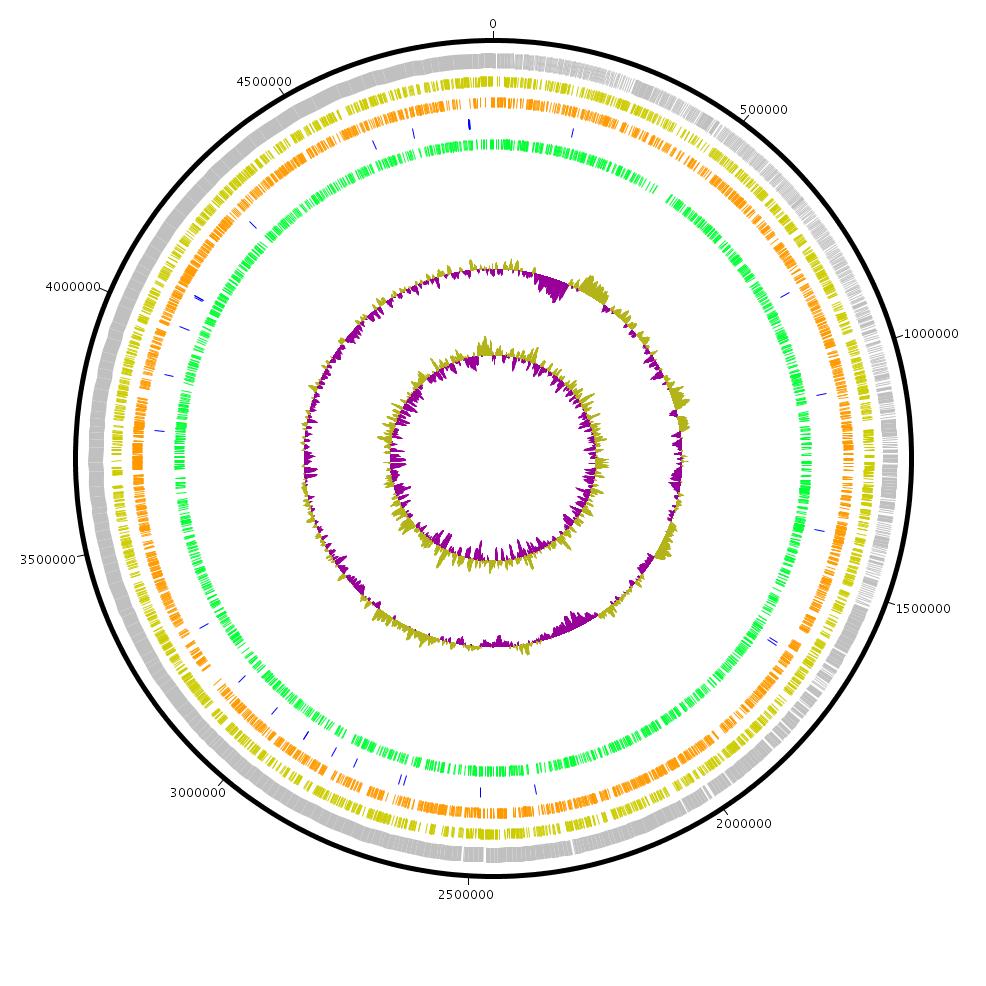

4.9 Mbp draft

*Segetibacter* genome

**B**

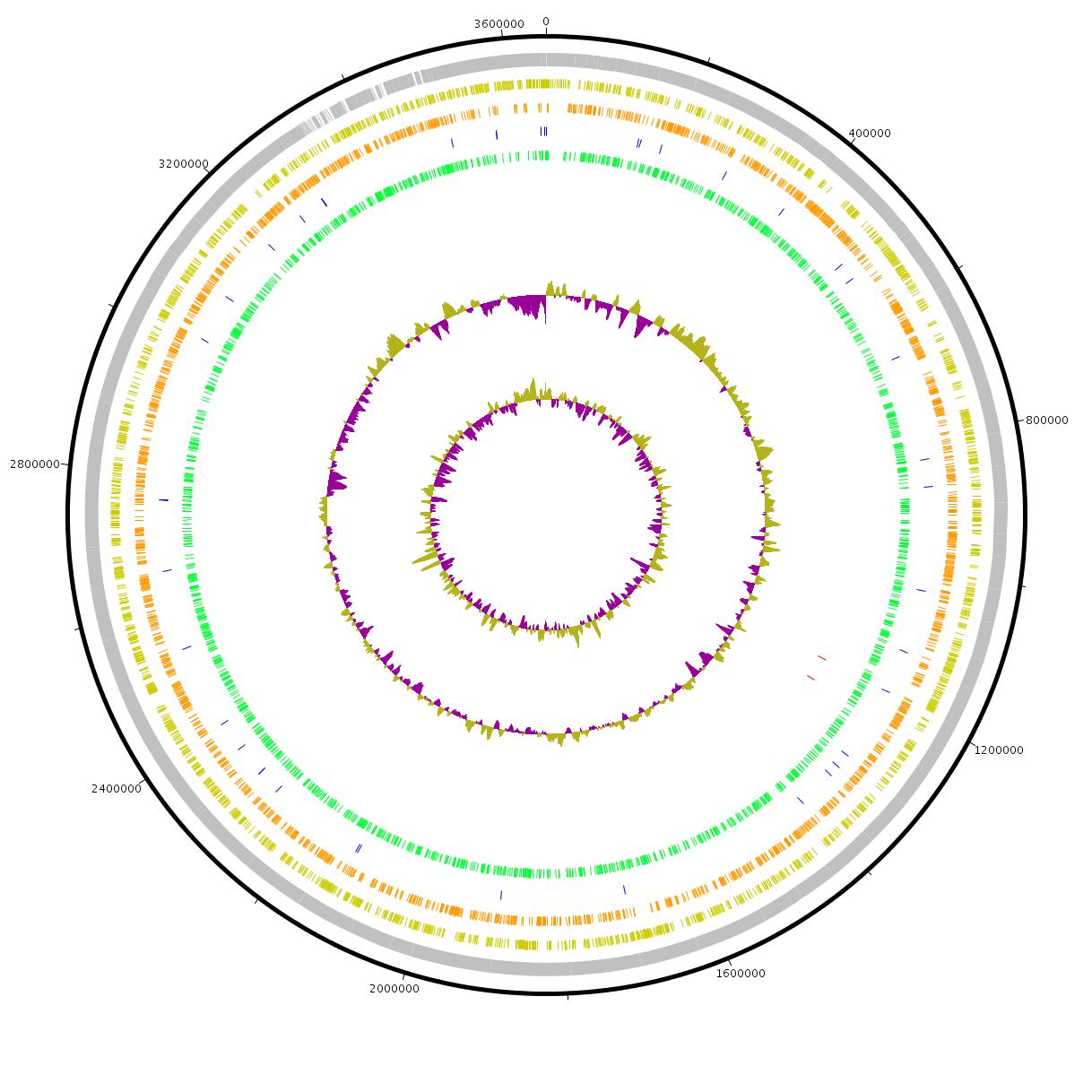

3.7 Mbp draft

*Pyrinomonas* genome

**Supplementary Figure 4.** Isotope bi-plot for organic materials of Mackay Glacier soils. The values plotted are mean δ^13^C and δ^15^N values.

δ15N / ‰

| δ13C / ‰ | 15N corr |
| --- | --- |
| -31.3442 | -10.3639 |
| -26.4212 | -9.75532 |
| -27.3787 | -0.406133 |
| -28.4635 | -8.08381 |
| -29.9548 | -5.78624 |
| -26.4955 | -2.31412 |
| -28.3478 | -1.71269 |
| -26.4724 | 0.0186398 |
| -28.1404 | -6.6047 |
| -26.529 | -5.02003 |
| -26.8151 | -2.40097 |
| -24.8443 | 2.72048 |
| -28.6856 | -8.24733 |
| -23.7179 | -6.89108 |
| -23.4485 | -0.792546 |
| -27.0079 | -11.5826 |
| -27.5508 | -13.7787 |
| -28.9109 | -11.7775 |
| -27.2912 | -8.67042 |

**Supplementary Table 1.** The soil physicochemical and environmental characteristics of the 18 sites along the Mackay Glacier ecotone.

| **Sample ID** | **Location Name** | | **Latitude** | | **Longitude** | | **Soil pH** | **Altitude (m.a.s.)** | | **Stone Fraction (%)** |
| --- | --- | --- | --- | --- | --- | --- | --- | --- | --- | --- |
| BG12-3 | Mount Benson | | 76° 52.144' S | | 161° 45.148' E | | 8.20 | 1109 | | 4 |
| CN-4 | Cliff Nunatak | | 76° 06.627' S | | 162° 00.839' E | | 7.95 | 220 | | 48 |
| FI-1 | Flatiron | | 77° 00.377' S | | 162° 23.841' E | | 8.35 | 517 | | 2 |
| MG3-2 | Mackay Glacier | | 76° 47.005' S | | 161° 27.014' E | | 7.53 | 189 | | 1 |
| MG6-4 | Mackay Glacier | | 76° 49.358' S | | 162° 06.422' E | | 7.86 | 763 | | 11 |
| MGM-3 | Mount George Murray | | 76° 09.646' S | | 162° 00.950' E | | 7.65 | 157 | | 38 |
| MS1-1 | Mount Seuss | | 77° 01.149' S | | 161° 46.931' E | | 7.58 | 556 | | 2 |
| MS2-2 | Mount Seuss | | 77° 00.654' S | | 161° 46.033' E | | 7.79 | 497 | | 2 |
| MS3-5 | Mount Seuss | | 77° 01.091' S | | 161° 44.507' E | | 7.81 | 547 | | 1 |
| MS4-1 | Mount Seuss | | 77° 01.384' S | | 161° 42.794' E | | 7.74 | 527 | | 1 |
| MS5-1 | Mount Seuss | | 77° 01.572' S | | 161° 40.490' E | | 7.77 | 576 | | 1 |
| MS6-5 | Mount Seuss | | 77° 01.102' S | | 161° 42.587' E | | 8.70 | 488 | | 30 |
| MS7-5 | Mount Seuss | | 77° 01.482' S | | 161° 45.550' E | | 7.94 | 440 | | 34 |
| MtG-4 | Mount Gran | | 76° 57.133' S | | 161° 24.263' E | | 8.17 | 652 | | 31 |
| MtG22-5 | Mount Gran | | 76° 58.027' S | | 161° 09.730' E | | 7.94 | 963 | | 2 |
| PT-2 | Pegtop Mountain | | 77° 02.762' S | | 161° 21.559' E | | 8.21 | 811 | | 2 |
| TG1-5 | Towle Glacier | | 76° 39.277' S | | 161° 05.577' E | | 8.14 | 1017 | | 1 |
| TG5-1 | Towle Glacier | | 76° 43.731' S | | 161° 00.699' E | | 8.08 | 729 | | 3 |
| **Sample ID** | **Mg (cmol(+)/kg)** | **Ca (cmol(+)/kg)** | | **K (mg/kg)** | | **Na (cmol(+)/kg)** | | | **Soil Respiration (µg C/g^-1^ soil/d^-1^)** | |
| BG12-3 | 0.89 | 2.16 | | 61 | | 0.60 | | | 0.153 | |
| CN-4 | 0.32 | 0.48 | | 26 | | 0.07 | | | 0.401 | |
| FI-1 | 0.99 | 10.3 | | 299 | | 1.95 | | | 0.916 | |
| MG3-2 | 0.78 | 1.29 | | 35 | | 0.13 | | | 0.508 | |
| MG6-4 | 0.63 | 1.49 | | 83 | | 0.33 | | | 0.212 | |
| MGM-3 | 0.47 | 1.00 | | 32 | | 0.34 | | | 0.231 | |
| MS1-1 | 2.10 | 12.8 | | 116 | | 1.92 | | | 0.403 | |
| MS2-2 | 1.41 | 6.33 | | 78 | | 1.40 | | | 0.139 | |
| MS3-5 | 1.33 | 6.04 | | 58 | | 0.70 | | | 0.149 | |
| MS4-1 | 1.02 | 6.43 | | 56 | | 0.37 | | | 0.075 | |
| MS5-1 | 0.98 | 4.02 | | 54 | | 0.49 | | | 0.801 | |
| MS6-5 | 0.91 | 4.95 | | 44 | | 0.94 | | | 0.171 | |
| MS7-5 | 2.15 | 7.97 | | 87 | | 0.53 | | | 0.346 | |
| MtG-4 | 0.44 | 3.70 | | 22 | | 0.13 | | | 0.295 | |
| MtG22-5 | 1.37 | 4.48 | | 75 | | 0.32 | | | 0.934 | |
| PT-2 | 0.53 | 1.09 | | 15 | | 0.07 | | | 0.938 | |
| TG1-5 | 1.62 | 2.80 | | 29 | | 0.07 | | | 0.091 | |
| TG5-1 | 1.57 | 2.97 | | 27 | | 0.28 | | | 0.566 | |

**Supplementary Table 2.** Respiration rates (soil CO_2_ release).

| **Sample ID** | **Average rate (μg/C soil/h^-1^)** | **S.E.** |
| --- | --- | --- |
| BG12-3 | 0.153 |  |
| CN-4 | 0.401 |  |
| F1-1 | 0.916 | 0.633 |
| MG3-2 | 0.508 |  |
| MG6-4 | 0.212 | 0.116 |
| MGM-3 | 0.231 | 0.058 |
| MS1-1 | 0.403 |  |
| MS2-2 | 0.139 |  |
| MS3-5 | 0.149 |  |
| MS4-1 | 0.075 |  |
| MS5-1 | 0.801 | 0.638 |
| MS6-5 | 0.171 | 0.012 |
| MS7-5 | 0.346 | 0.133 |
| MtG22-5 | 0.934 | 0.653 |
| MtG-4 | 0.295 | 0.044 |
| PT-2 | 0.938 | 0.638 |
| TG1-5 | 0.091 |  |
| TG5-1 | 0.566 | 0.130 |

**Supplementary Table 3.** Soil metagenome sequencing statistics.

^a^ QC – Quality Control; bp – base pairs

| **Sample ID** | **Pre-QC no. of reads^a^** | **Merged reads** | **Average read length (bp)** | **Mean G+C content (%)** | ***N_25_/N_50_/N_75_* (bp)** |
| --- | --- | --- | --- | --- | --- |
| BG12-3 | 10,336,306 | 7,730,414 | 304.46 | 48.04 | 403/340/276 |
| CN-4 | 12,152,688 | 9,464,484 | 304.96 | 55.08 | 404/343/280 |
| FI-1 | 21,825,223 | 18,261,157 | 281.25 | 47.36 | 387/320/256 |
| MG3-2 | 9,346,257 | 6,407,139 | 320.24 | 47.56 | 417/360/297 |
| MG6-4 | 10,197,250 | 8,126,681 | 299.41 | 49.64 | 400/338/276 |
| MGM-3 | 12,034,349 | 9,469,426 | 294.87 | 50.45 | 395/332/271 |
| MS1-1 | 11,236,067 | 8,206,577 | 310.64 | 52.98 | 408/347/285 |
| MS2-2 | 11,530,683 | 8,846,170 | 302.59 | 60.17 | 402/341/279 |
| MS3-5 | 13,036,230 | 10,765,138 | 291.02 | 48.58 | 394/330/267 |
| MS4-1 | 13,696,440 | 11,178,757 | 287.54 | 50.42 | 389/323/260 |
| MS5-1 | 11,535,330 | 8,850,331 | 309.99 | 53.94 | 408/350/288 |
| MS6-5 | 11,891,994 | 9,246,318 | 304.69 | 53.97 | 405/345/282 |
| MS7-5 | 12,767,348 | 10,609,771 | 287.46 | 58.68 | 391/328/265 |
| MtG-4 | 11,026,310 | 8,204,285 | 308.91 | 45.71 | 409/349/286 |
| MtG22-5 | 15,138,920 | 10,510,936 | 313.48 | 60.24 | 411/354/293 |
| PT-2 | 10,888,908 | 8,284,819 | 308.24 | 47.68 | 407/348/285 |
| TG1-5 | 7,640,231 | 4,023,268 | 285.60 | 53.57 | 382/331/270 |
| TG5-1 | 7,371,647 | 3,980,751 | 218.44 | 46.93 | 380/327/264 |

**Supplementary Table 4.** The relative abundance of major phyla across the 18 soils in arranged in order of increasing altitude. The numbers in cells indicate percentage abundance. The cell colours are based on relative abundances with the highest values in red.

| **Domain** | **Phylum** | **MGM-3** | **MG3-2** | **CN-4** | **MS7-5** | **MS6-5** | **MS2-2** | **FI-1** | **MS4-1** | **MS3-5** | **MS1-1** | **MS5-1** | **MtG-4** | **TG5-1** | **MG6-4** | **PT-2** | **MtG22-5** | **TG1-5** | **BG12-3** |
| --- | --- | --- | --- | --- | --- | --- | --- | --- | --- | --- | --- | --- | --- | --- | --- | --- | --- | --- | --- |
| Archaea | *Crenarchaeota* | 0,04 | 0,00 | 0,00 | 0,00 | 0,02 | 0,00 | 0,00 | 0,02 | 0,00 | 0,03 | 0,02 | 0,01 | 0,00 | 0,00 | 0,01 | 0,08 | 0,00 | 0,04 |
|  | *Euryarchaeota* | 0,19 | 0,49 | 0,25 | 0,38 | 0,27 | 0,31 | 0,38 | 0,34 | 0,39 | 0,27 | 0,27 | 0,27 | 0,34 | 0,26 | 0,31 | 0,25 | 0,24 | 0,28 |
|  | *Thaumarchaeota* | 0,50 | 0,08 | 0,07 | 0,33 | 0,32 | 0,08 | 0,29 | 0,55 | 0,18 | 0,52 | 0,41 | 0,13 | 0,12 | 0,11 | 0,16 | 1,11 | 0,03 | 1,15 |
|  | Unclassified archaea | 0,00 | 0,00 | 0,00 | 0,00 | 0,00 | 0,00 | 0,00 | 0,02 | 0,00 | 0,02 | 0,01 | 0,01 | 0,00 | 0,00 | 0,02 | 0,00 | 0,00 | 0,00 |
| Bacteria | *Acidobacteria* | 8,68 | 19,38 | 20,20 | 16,37 | 21,84 | 19,13 | 4,07 | 18,98 | 25,03 | 13,93 | 15,54 | 16,82 | 25,99 | 18,34 | 22,52 | 8,73 | 25,30 | 15,41 |
|  | *Actinobacteria* | 0,93 | 3,41 | 2,44 | 4,93 | 1,41 | 8,85 | 2,82 | 1,92 | 3,07 | 2,35 | 2,01 | 1,23 | 1,79 | 1,18 | 2,90 | 3,70 | 1,53 | 1,86 |
|  | *Aquificae* | 0,02 | 0,00 | 0,00 | 0,00 | 0,03 | 0,00 | 0,00 | 0,03 | 0,00 | 0,00 | 0,02 | 0,03 | 0,04 | 0,03 | 0,03 | 0,00 | 0,02 | 0,00 |
|  | *Armatimonadetes* | 0,09 | 0,23 | 0,71 | 0,11 | 0,14 | 0,18 | 0,17 | 0,12 | 0,13 | 0,08 | 0,12 | 0,10 | 0,13 | 0,09 | 0,34 | 0,06 | 0,13 | 0,09 |
|  | *Bacteroidetes* | 65,98 | 34,46 | 39,34 | 50,99 | 51,73 | 35,26 | 52,31 | 52,87 | 37,14 | 58,93 | 47,49 | 57,48 | 46,28 | 59,43 | 44,76 | 72,64 | 52,94 | 60,48 |
|  | *Chloroflexi* | 0,54 | 2,36 | 4,12 | 1,95 | 0,85 | 1,77 | 1,28 | 1,11 | 1,14 | 0,96 | 1,37 | 1,23 | 1,48 | 0,66 | 1,28 | 0,55 | 0,94 | 0,72 |
|  | *Cyanobacteria* | 8,86 | 7,82 | 8,76 | 3,71 | 5,17 | 7,83 | 3,84 | 4,96 | 8,44 | 8,65 | 6,00 | 3,11 | 4,22 | 5,78 | 9,59 | 1,82 | 3,47 | 3,73 |
|  | *Deinococcus-Thermus* | 0,21 | 1,00 | 0,35 | 0,60 | 0,36 | 1,12 | 1,71 | 0,37 | 0,75 | 0,64 | 0,36 | 0,21 | 0,35 | 0,45 | 0,45 | 0,17 | 0,25 | 0,46 |
|  | *Firmicutes* | 1,23 | 4,04 | 2,03 | 2,27 | 2,02 | 2,59 | 6,17 | 2,39 | 3,32 | 1,94 | 1,80 | 1,96 | 2,53 | 1,96 | 2,37 | 1,33 | 1,83 | 2,24 |
|  | *Fusobacteria* | 0,02 | 0,00 | 0,00 | 0,00 | 0,00 | 0,00 | 0,00 | 0,02 | 0,00 | 0,00 | 0,01 | 0,02 | 0,00 | 0,02 | 0,02 | 0,00 | 0,00 | 0,00 |
|  | *Gemmatimonadetes* | 0,30 | 0,50 | 0,23 | 0,88 | 0,57 | 0,62 | 0,89 | 0,78 | 0,35 | 0,68 | 2,10 | 0,64 | 1,17 | 0,17 | 0,26 | 0,26 | 0,21 | 0,19 |
|  | *Nitrospinae* | 0,01 | 0,00 | 0,00 | 0,00 | 0,00 | 0,00 | 0,00 | 0,01 | 0,00 | 0,00 | 0,02 | 0,02 | 0,02 | 0,00 | 0,02 | 0,00 | 0,00 | 0,00 |
|  | *Nitrospirae* | 0,16 | 0,11 | 0,05 | 0,36 | 0,13 | 0,20 | 0,10 | 0,13 | 0,11 | 0,10 | 0,36 | 0,12 | 0,11 | 0,05 | 0,08 | 0,10 | 0,05 | 0,09 |
|  | *Planctomycetes* | 0,61 | 1,65 | 1,02 | 0,83 | 0,74 | 1,23 | 1,82 | 0,86 | 1,18 | 0,66 | 1,12 | 0,75 | 0,84 | 0,67 | 0,86 | 0,52 | 0,77 | 0,70 |
|  | *Proteobacteria* | 7,88 | 18,46 | 8,90 | 13,00 | 10,09 | 16,76 | 17,14 | 9,58 | 13,97 | 7,40 | 13,52 | 9,15 | 10,24 | 7,87 | 8,75 | 6,20 | 7,45 | 9,09 |
|  | *Spirochaetes* | 0,14 | 0,28 | 0,12 | 0,10 | 0,17 | 0,13 | 0,26 | 0,20 | 0,23 | 0,15 | 0,15 | 0,20 | 0,17 | 0,15 | 0,16 | 0,13 | 0,15 | 0,19 |
|  | *Synergistetes* | 0,00 | 0,00 | 0,00 | 0,00 | 0,00 | 0,00 | 0,00 | 0,01 | 0,00 | 0,00 | 0,01 | 0,01 | 0,00 | 0,00 | 0,01 | 0,00 | 0,00 | 0,00 |
|  | *Thermotogae* | 0,02 | 0,00 | 0,02 | 0,00 | 0,02 | 0,00 | 0,00 | 0,02 | 0,00 | 0,00 | 0,02 | 0,02 | 0,02 | 0,02 | 0,02 | 0,00 | 0,00 | 0,00 |
|  | Unclassified bacteria | 1,86 | 3,14 | 2,77 | 2,11 | 2,48 | 2,22 | 1,39 | 2,35 | 2,70 | 1,74 | 3,17 | 3,17 | 2,67 | 2,01 | 2,41 | 1,57 | 2,63 | 2,23 |
|  | *Verrucomicrobia* | 1,44 | 1,49 | 3,21 | 0,80 | 1,35 | 1,30 | 4,69 | 1,52 | 1,13 | 0,67 | 3,80 | 2,73 | 1,17 | 0,44 | 2,28 | 0,52 | 1,80 | 0,52 |
| Eukaryota | *Alveolata* | 0,00 | 0,00 | 0,04 | 0,00 | 0,00 | 0,00 | 0,00 | 0,00 | 0,00 | 0,00 | 0,02 | 0,02 | 0,00 | 0,00 | 0,01 | 0,00 | 0,00 | 0,00 |
|  | *Amoebozoa* | 0,03 | 0,11 | 0,00 | 0,00 | 0,02 | 0,00 | 0,00 | 0,14 | 0,00 | 0,00 | 0,02 | 0,14 | 0,00 | 0,00 | 0,01 | 0,06 | 0,00 | 0,00 |
|  | *Opisthokonta* | 0,18 | 0,84 | 4,89 | 0,17 | 0,19 | 0,29 | 0,54 | 0,55 | 0,75 | 0,22 | 0,16 | 0,29 | 0,24 | 0,23 | 0,29 | 0,12 | 0,23 | 0,48 |
|  | *Stramenopiles* | 0,03 | 0,00 | 0,00 | 0,00 | 0,00 | 0,00 | 0,11 | 0,02 | 0,00 | 0,00 | 0,01 | 0,02 | 0,00 | 0,00 | 0,00 | 0,00 | 0,00 | 0,00 |
|  | *Viridiplantae* | 0,06 | 0,09 | 0,46 | 0,00 | 0,05 | 0,05 | 0,00 | 0,09 | 0,00 | 0,03 | 0,04 | 0,08 | 0,03 | 0,03 | 0,03 | 0,03 | 0,03 | 0,00 |
| Viruses | Caudovirales | 0,01 | 0,07 | 0,02 | 0,12 | 0,03 | 0,09 | 0,00 | 0,04 | 0,00 | 0,04 | 0,04 | 0,04 | 0,06 | 0,03 | 0,06 | 0,05 | 0,00 | 0,04 |

**Supplementary Table 5.** Functional genes analysed for carbon and nitrogen cycling.

| **Cycle** | **Step** | **Gene** | **KO Terms** |
| --- | --- | --- | --- |
| **Carbon** | Aerobic Carbon Fixation | phosphoribulokinase (prk*B*) | K00855 |
|  |  | Ribulose-1,5-bisphosphate carboxylase oxygenase (RuBisCO) large chain (rbc*L*) | K01601 |
|  |  | Ribulose-1,5-bisphosphate carboxylase oxygenase (RuBisCO) small chain (rbc*S*) | K01602 |
|  | Aerobic Respiration | cytochrome c oxidase, aa_3_-type subunit I (cox*I*) | K02256 |
|  |  | cytochrome c oxidase, aa_3_-type subunit III (cox*III*) | K02262 |
|  |  | cytochrome c oxidase, aa_3_-type subunit I (cox*A*) | K02274 |
|  |  | cytochrome c oxidase, aa_3_-type subunit III (cox*C*) | K02276 |
|  | Anaerobic Carbon Fixation | 2-oxogluterate:ferredoxin oxidoreductase subunit alpha (kor*A*) | K00174 |
|  |  | 2-oxogluterate:ferredoxin oxidoreductase subunit beta (kor*B*) | K00175 |
|  |  | fumarate reductase flavoprotein subunit (frd*A*) | K00244 |
|  |  | adenosinetriphosphate (ATP) citrate lyase (ACLY) | K01648 |
|  | Carbon monoxide oxidation | carbon-monoxide dehydrogenase small subunit (cox*S*) | K03518 |
|  |  | carbon-monoxide dehydrogenase medium subunit (cut*M*, cox*M*) | K03519 |
|  |  | carbon-monoxide dehydrogenase large subunit (cut*L*, cox*L*) | K03520 |
|  | Fermentation | L-lactate dehydrogenase (ldh) | K00016 |
|  | Methanogenesis | coenzyme M methyl reductase beta subunit (mcr*B*) | K00400 |
| **Nitrogen** | Ammonification | formate-dependent nitrite reductase periplasmic cytochrome c552 (nrf*A*) | K03385 |
|  | Anammox (SRAO) | hydroxylamine oxidoreductase/hydrazine oxidoreductase (hao/hzo) | K10535 |
|  | Denitrification | nitrous oxide reductase (nos*Z*) | K00376 |
|  |  | nitric-oxide reductase (nor*C*) | K02305 |
|  |  | nitric-oxide reductase (nor*B*) | K04561 |
|  | Nitrate Reduction | periplasmic nitrate reductase (nap*A*) | K02567 |
|  |  | cytochrome c-type protein (nap*B*) | K02568 |
|  | Nitrate Reduction & Nitrite oxidation | nitrate reductase alpha & nitrite oxidoreductase (nar*G*/nxr*A*) | K00370 |
|  |  | nitrate reductase beta & nitrite oxidoreductase (nar*H*/nxr*B*) | K00371 |
|  | Nitrification | ammonia monooxygenase subunit A (amo*A*, pmo*A*) | K10944 |
|  |  | ammonia monooxygenase subunit B (amo*B*, pmo*B*) | K10945 |
|  |  | ammonia monooxygenase subunit C (amo*C*, pmo*C*) | K10946 |
|  | Nitrogen Assimilation | glutamate synthase (NADPH/NADH) large chain (glt*B*) | K00265 |
|  |  | glutamate synthase (ferredoxin-dependent) (glt*S*) | K00284 |
|  |  | assimilatory nitrate reductase (nas) | K00360 |
|  |  | glutamine synthetase (gln*A*) | K01915 |
|  | Nitrogen Fixation | nitrogenase molybdenum-iron protein alpha chain (nif*D*) | K02586 |
|  |  | nitrogenase iron protein (nif*H*) | K02588 |
|  |  | nitrogenase molybdenum-iron protein beta chain (nif*K*) | K02591 |
|  | Nitrogen Mineralization | glutamate dehydrogenase (gdh*A*) | K00260 |

**Supplementary Table 6.** Taxonomic assignments of functional genes involved in C and N turnover from each metagenome.

| **Gene** | **Phylum** | **BG12-3** | **CN-4** | **FI-1** | **MG3-2** | **MG6-4** | **MGM-3** | **MS1-1** | **MS2-2** | **MS3-5** | **MS4-1** | **MS5-1** | **MS6-5** | **MS7-5** | **MtG-4** | **MtG22-5** | **PT-2** | **TG1-5** | **TG5-1** |
| --- | --- | --- | --- | --- | --- | --- | --- | --- | --- | --- | --- | --- | --- | --- | --- | --- | --- | --- | --- |
| ACLY | Other | 0 | 8 | 0 | 0 | 0 | 0 | 0 | 0 | 0 | 0 | 1 | 0 | 0 | 3 | 0 | 0 | 0 | 0 |
| amo*ABC* | *Bacteroidetes* | 0 | 0 | 0 | 0 | 0 | 0 | 0 | 0 | 0 | 0 | 2 | 0 | 0 | 0 | 0 | 0 | 0 | 0 |
| amo*ABC* | *Cyanobacteria* | 0 | 0 | 0 | 0 | 0 | 0 | 0 | 0 | 0 | 0 | 1 | 0 | 0 | 0 | 0 | 0 | 0 | 0 |
| amo*ABC* | *Proteobacteria* | 0 | 0 | 0 | 0 | 0 | 0 | 0 | 0 | 0 | 0 | 0 | 0 | 0 | 1 | 0 | 0 | 0 | 0 |
| amo*ABC* | Unclassified Archaea | 2 | 1 | 0 | 0 | 1 | 2 | 1 | 0 | 0 | 3 | 0 | 0 | 0 | 0 | 1 | 1 | 0 | 0 |
| CDO1 | *Acidobacteria* | 7 | 8 | 1 | 1 | 13 | 7 | 9 | 9 | 4 | 12 | 16 | 18 | 8 | 30 | 1 | 25 | 12 | 16 |
| CDO1 | Other | 0 | 0 | 0 | 0 | 0 | 0 | 0 | 0 | 0 | 0 | 0 | 0 | 0 | 0 | 0 | 1 | 0 | 0 |
| cox*AC* | *Acidobacteria* | 16 | 50 | 2 | 10 | 30 | 40 | 31 | 13 | 14 | 65 | 36 | 36 | 18 | 83 | 18 | 56 | 66 | 42 |
| cox*AC* | *Actinobacteria* | 0 | 0 | 1 | 0 | 0 | 0 | 1 | 4 | 0 | 2 | 3 | 0 | 0 | 1 | 7 | 6 | 1 | 0 |
| cox*AC* | *Bacteroidetes* | 31 | 46 | 10 | 8 | 48 | 104 | 36 | 12 | 7 | 59 | 80 | 35 | 14 | 110 | 43 | 55 | 46 | 46 |
| cox*AC* | *Chloroflexi* | 3 | 8 | 0 | 2 | 9 | 8 | 5 | 2 | 3 | 7 | 12 | 7 | 3 | 12 | 2 | 11 | 10 | 7 |
| cox*AC* | *Cyanobacteria* | 0 | 15 | 0 | 0 | 1 | 25 | 5 | 1 | 0 | 5 | 2 | 1 | 2 | 2 | 0 | 17 | 0 | 0 |
| cox*AC* | *Gemmatimonadetes* | 0 | 1 | 1 | 1 | 0 | 1 | 0 | 1 | 0 | 3 | 0 | 1 | 2 | 5 | 0 | 0 | 0 | 6 |
| cox*AC* | Other | 0 | 10 | 9 | 0 | 1 | 1 | 0 | 1 | 0 | 0 | 2 | 0 | 1 | 3 | 2 | 1 | 1 | 0 |
| cox*AC* | *Planctomycetes* | 1 | 2 | 0 | 0 | 5 | 2 | 2 | 2 | 1 | 6 | 0 | 5 | 0 | 14 | 0 | 8 | 4 | 10 |
| cox*AC* | *Proteobacteria* | 1 | 4 | 3 | 0 | 2 | 6 | 2 | 2 | 3 | 5 | 0 | 7 | 1 | 20 | 3 | 1 | 1 | 6 |
| cox*AC* | Unclassified Archaea | 1 | 0 | 0 | 0 | 0 | 3 | 0 | 0 | 0 | 2 | 0 | 0 | 0 | 0 | 2 | 0 | 0 | 0 |
| cox*AC* | Unclassified Bacteria | 22 | 61 | 0 | 17 | 54 | 57 | 51 | 33 | 22 | 101 | 70 | 70 | 26 | 134 | 24 | 86 | 62 | 88 |
| cox*AC* | *Verrucomicrobia* | 0 | 6 | 0 | 0 | 0 | 9 | 1 | 1 | 1 | 8 | 15 | 7 | 4 | 9 | 0 | 20 | 8 | 6 |
| cox*I* - cox*III* | Other | 0 | 2 | 1 | 3 | 0 | 0 | 0 | 0 | 0 | 0 | 0 | 0 | 0 | 2 | 0 | 0 | 0 | 0 |
| cox*I* - cox*III* | *Proteobacteria* | 0 | 0 | 0 | 0 | 0 | 0 | 0 | 0 | 0 | 1 | 0 | 0 | 0 | 0 | 0 | 0 | 0 | 0 |
| cox*LMS* | *Acidobacteria* | 0 | 5 | 0 | 0 | 0 | 5 | 1 | 2 | 0 | 17 | 11 | 0 | 3 | 10 | 4 | 1 | 3 | 0 |
| cox*LMS* | *Actinobacteria* | 0 | 0 | 0 | 0 | 0 | 0 | 3 | 0 | 0 | 0 | 0 | 0 | 2 | 0 | 1 | 1 | 1 | 0 |
| cox*LMS* | *Bacteroidetes* | 6 | 0 | 0 | 0 | 17 | 40 | 12 | 2 | 0 | 21 | 16 | 18 | 2 | 54 | 4 | 9 | 9 | 8 |
| cox*LMS* | *Chloroflexi* | 0 | 2 | 0 | 0 | 0 | 3 | 0 | 1 | 1 | 0 | 0 | 0 | 0 | 2 | 0 | 1 | 0 | 1 |
| cox*LMS* | *Cyanobacteria* | 0 | 1 | 0 | 0 | 0 | 0 | 0 | 0 | 0 | 0 | 0 | 0 | 0 | 0 | 0 | 0 | 0 | 0 |
| cox*LMS* | *Gemmatimonadetes* | 0 | 0 | 0 | 1 | 0 | 0 | 0 | 0 | 0 | 0 | 0 | 0 | 0 | 0 | 0 | 0 | 0 | 1 |
| cox*LMS* | Other | 0 | 1 | 2 | 0 | 0 | 1 | 0 | 0 | 0 | 1 | 1 | 0 | 0 | 3 | 0 | 3 | 3 | 0 |
| cox*LMS* | *Proteobacteria* | 0 | 1 | 1 | 0 | 0 | 1 | 0 | 0 | 0 | 2 | 0 | 0 | 0 | 0 | 0 | 0 | 0 | 0 |
| cox*LMS* | Unclassified Bacteria | 1 | 9 | 0 | 0 | 7 | 10 | 4 | 3 | 2 | 7 | 22 | 5 | 1 | 15 | 5 | 5 | 5 | 3 |
| cys*C* | *Acidobacteria* | 0 | 2 | 0 | 0 | 0 | 1 | 0 | 0 | 0 | 3 | 1 | 0 | 0 | 0 | 0 | 0 | 0 | 0 |
| cys*C* | *Actinobacteria* | 0 | 0 | 0 | 0 | 2 | 1 | 2 | 2 | 0 | 1 | 0 | 0 | 1 | 1 | 0 | 1 | 1 | 1 |
| cys*C* | *Bacteroidetes* | 1 | 0 | 0 | 0 | 0 | 2 | 6 | 2 | 1 | 5 | 7 | 3 | 1 | 9 | 1 | 1 | 1 | 1 |
| cys*C* | *Chloroflexi* | 0 | 1 | 0 | 0 | 0 | 0 | 0 | 0 | 0 | 0 | 0 | 0 | 0 | 0 | 0 | 0 | 0 | 0 |
| cys*C* | *Cyanobacteria* | 0 | 0 | 0 | 0 | 0 | 1 | 1 | 1 | 0 | 0 | 0 | 0 | 0 | 0 | 0 | 3 | 0 | 0 |
| cys*C* | *Gemmatimonadetes* | 0 | 0 | 0 | 0 | 0 | 0 | 0 | 0 | 0 | 0 | 0 | 0 | 0 | 1 | 0 | 0 | 0 | 0 |
| cys*C* | Other | 0 | 1 | 1 | 0 | 0 | 1 | 0 | 0 | 0 | 2 | 1 | 1 | 0 | 0 | 0 | 6 | 0 | 1 |
| cys*C* | *Proteobacteria* | 0 | 1 | 0 | 0 | 0 | 0 | 0 | 0 | 0 | 1 | 0 | 0 | 0 | 1 | 0 | 0 | 0 | 1 |
| cys*C* | Unclassified Bacteria | 0 | 4 | 0 | 0 | 4 | 8 | 7 | 1 | 1 | 6 | 9 | 1 | 1 | 9 | 2 | 7 | 3 | 1 |
| cys*DN* | *Acidobacteria* | 0 | 2 | 0 | 0 | 0 | 0 | 0 | 0 | 0 | 0 | 0 | 0 | 0 | 0 | 0 | 1 | 0 | 0 |
| cys*DN* | *Actinobacteria* | 0 | 1 | 1 | 0 | 0 | 0 | 1 | 1 | 0 | 0 | 1 | 0 | 2 | 0 | 0 | 0 | 0 | 0 |
| cys*DN* | *Bacteroidetes* | 26 | 21 | 4 | 6 | 36 | 103 | 31 | 10 | 5 | 55 | 45 | 27 | 16 | 115 | 57 | 42 | 31 | 42 |
| cys*DN* | Other | 1 | 0 | 1 | 0 | 0 | 1 | 0 | 0 | 0 | 0 | 0 | 0 | 0 | 0 | 0 | 0 | 0 | 0 |
| cys*DN* | *Proteobacteria* | 1 | 1 | 0 | 0 | 0 | 2 | 1 | 3 | 0 | 0 | 0 | 3 | 2 | 1 | 0 | 0 | 0 | 0 |
| cys*DN* | Unclassified Bacteria | 3 | 4 | 0 | 0 | 2 | 6 | 7 | 0 | 2 | 2 | 6 | 7 | 2 | 20 | 2 | 12 | 2 | 6 |
| cys*DN* | *Verrucomicrobia* | 0 | 0 | 0 | 0 | 0 | 0 | 0 | 1 | 0 | 0 | 1 | 0 | 0 | 3 | 0 | 3 | 1 | 0 |
| frd*A* | *Actinobacteria* | 0 | 0 | 1 | 0 | 0 | 0 | 0 | 0 | 0 | 0 | 0 | 0 | 0 | 0 | 0 | 0 | 0 | 0 |
| frd*A* | Other | 0 | 0 | 0 | 0 | 0 | 0 | 0 | 0 | 0 | 0 | 0 | 0 | 1 | 0 | 0 | 0 | 0 | 0 |
| frd*A* | *Proteobacteria* | 0 | 0 | 0 | 0 | 0 | 0 | 0 | 0 | 0 | 1 | 0 | 0 | 0 | 0 | 0 | 0 | 0 | 0 |
| frd*A* | Unclassified Bacteria | 0 | 0 | 0 | 0 | 0 | 1 | 0 | 0 | 0 | 0 | 0 | 0 | 0 | 0 | 0 | 1 | 0 | 0 |
| gdh*A* | *Acidobacteria* | 20 | 19 | 1 | 9 | 18 | 12 | 12 | 7 | 13 | 42 | 33 | 21 | 7 | 37 | 7 | 54 | 37 | 25 |
| gdh*A* | *Actinobacteria* | 0 | 0 | 0 | 0 | 0 | 0 | 0 | 2 | 0 | 0 | 0 | 0 | 0 | 0 | 0 | 0 | 0 | 0 |
| gdh*A* | *Bacteroidetes* | 21 | 21 | 4 | 5 | 34 | 71 | 36 | 5 | 3 | 41 | 50 | 40 | 16 | 65 | 29 | 31 | 27 | 27 |
| gdh*A* | *Chloroflexi* | 0 | 0 | 0 | 0 | 0 | 0 | 2 | 0 | 0 | 1 | 1 | 1 | 1 | 3 | 0 | 2 | 1 | 0 |
| gdh*A* | *Cyanobacteria* | 0 | 2 | 0 | 0 | 1 | 1 | 0 | 0 | 0 | 0 | 0 | 0 | 0 | 0 | 0 | 0 | 0 | 0 |
| gdh*A* | *Gemmatimonadetes* | 0 | 0 | 0 | 0 | 0 | 1 | 0 | 0 | 2 | 0 | 0 | 1 | 0 | 0 | 1 | 0 | 1 | 0 |
| gdh*A* | Other | 0 | 2 | 1 | 0 | 0 | 2 | 1 | 0 | 0 | 3 | 3 | 1 | 0 | 9 | 0 | 2 | 0 | 1 |
| gdh*A* | *Planctomycetes* | 0 | 1 | 2 | 0 | 0 | 0 | 0 | 0 | 0 | 0 | 0 | 0 | 0 | 0 | 0 | 0 | 0 | 0 |
| gdh*A* | *Proteobacteria* | 0 | 1 | 0 | 0 | 0 | 6 | 0 | 0 | 0 | 0 | 0 | 0 | 0 | 4 | 0 | 1 | 0 | 0 |
| gdh*A* | Unclassified Archaea | 3 | 0 | 0 | 0 | 0 | 3 | 2 | 1 | 0 | 2 | 0 | 1 | 0 | 3 | 1 | 0 | 1 | 0 |
| gdh*A* | Unclassified Bacteria | 0 | 4 | 0 | 0 | 1 | 2 | 2 | 2 | 1 | 5 | 9 | 2 | 1 | 6 | 1 | 1 | 1 | 3 |
| gdh*A* | *Verrucomicrobia* | 0 | 0 | 0 | 0 | 0 | 0 | 0 | 0 | 0 | 1 | 1 | 1 | 0 | 1 | 0 | 0 | 0 | 0 |
| gln*A* | *Acidobacteria* | 12 | 22 | 0 | 5 | 21 | 13 | 12 | 7 | 8 | 23 | 21 | 16 | 4 | 34 | 6 | 41 | 32 | 20 |
| gln*A* | *Actinobacteria* | 1 | 0 | 0 | 0 | 0 | 2 | 2 | 12 | 2 | 3 | 3 | 0 | 2 | 2 | 7 | 7 | 2 | 3 |
| gln*A* | *Bacteroidetes* | 35 | 25 | 16 | 12 | 38 | 125 | 43 | 13 | 8 | 78 | 67 | 33 | 16 | 150 | 45 | 49 | 50 | 50 |
| gln*A* | *Chloroflexi* | 0 | 7 | 1 | 0 | 0 | 0 | 2 | 2 | 0 | 2 | 7 | 0 | 0 | 1 | 0 | 1 | 0 | 6 |
| gln*A* | *Cyanobacteria* | 0 | 10 | 0 | 0 | 4 | 7 | 9 | 2 | 1 | 5 | 2 | 1 | 2 | 0 | 0 | 9 | 0 | 0 |
| gln*A* | *Gemmatimonadetes* | 0 | 1 | 0 | 1 | 0 | 2 | 0 | 0 | 0 | 2 | 0 | 2 | 1 | 3 | 0 | 0 | 0 | 2 |
| gln*A* | Other | 0 | 3 | 5 | 0 | 1 | 3 | 0 | 0 | 1 | 1 | 0 | 2 | 0 | 4 | 0 | 2 | 1 | 0 |
| gln*A* | *Planctomycetes* | 0 | 0 | 0 | 0 | 0 | 1 | 0 | 0 | 1 | 1 | 0 | 0 | 2 | 0 | 0 | 0 | 0 | 0 |
| gln*A* | *Proteobacteria* | 0 | 4 | 0 | 0 | 2 | 11 | 0 | 1 | 1 | 4 | 0 | 6 | 0 | 4 | 3 | 4 | 0 | 2 |
| gln*A* | Unclassified Archaea | 2 | 0 | 0 | 0 | 1 | 2 | 2 | 0 | 0 | 3 | 0 | 1 | 0 | 3 | 3 | 0 | 0 | 0 |
| gln*A* | Unclassified Bacteria | 7 | 15 | 0 | 0 | 7 | 12 | 17 | 5 | 3 | 20 | 25 | 9 | 3 | 19 | 7 | 26 | 10 | 11 |
| gln*A* | *Verrucomicrobia* | 0 | 2 | 0 | 0 | 0 | 2 | 0 | 0 | 0 | 4 | 2 | 1 | 0 | 3 | 0 | 4 | 1 | 0 |
| glt*B* | *Acidobacteria* | 0 | 2 | 0 | 0 | 0 | 0 | 0 | 2 | 1 | 0 | 1 | 0 | 0 | 0 | 0 | 1 | 0 | 0 |
| glt*B* | *Actinobacteria* | 1 | 1 | 1 | 0 | 1 | 1 | 2 | 1 | 1 | 0 | 2 | 1 | 1 | 0 | 2 | 5 | 0 | 1 |
| glt*B* | *Bacteroidetes* | 21 | 32 | 9 | 5 | 40 | 82 | 30 | 13 | 5 | 37 | 33 | 16 | 10 | 90 | 28 | 47 | 30 | 29 |
| glt*B* | *Chloroflexi* | 0 | 2 | 0 | 0 | 0 | 0 | 0 | 0 | 0 | 0 | 0 | 0 | 0 | 0 | 0 | 1 | 1 | 0 |
| glt*B* | *Cyanobacteria* | 0 | 1 | 0 | 0 | 0 | 0 | 0 | 0 | 0 | 0 | 1 | 0 | 0 | 0 | 0 | 0 | 0 | 0 |
| glt*B* | Other | 0 | 3 | 0 | 0 | 0 | 0 | 0 | 2 | 0 | 1 | 0 | 1 | 0 | 1 | 0 | 1 | 0 | 0 |
| glt*B* | *Planctomycetes* | 0 | 0 | 0 | 0 | 0 | 0 | 0 | 0 | 0 | 0 | 0 | 0 | 0 | 0 | 0 | 1 | 0 | 0 |
| glt*B* | *Proteobacteria* | 0 | 3 | 0 | 0 | 0 | 1 | 0 | 0 | 2 | 0 | 0 | 2 | 0 | 0 | 1 | 0 | 0 | 0 |
| glt*B* | Unclassified Bacteria | 1 | 2 | 0 | 0 | 0 | 2 | 1 | 2 | 1 | 1 | 3 | 1 | 2 | 2 | 1 | 6 | 0 | 2 |
| glt*B* | *Verrucomicrobia* | 0 | 0 | 0 | 0 | 0 | 1 | 0 | 0 | 0 | 0 | 0 | 0 | 0 | 0 | 0 | 0 | 0 | 0 |
| glt*S* | *Actinobacteria* | 0 | 0 | 0 | 0 | 0 | 1 | 0 | 0 | 0 | 0 | 0 | 0 | 0 | 0 | 0 | 0 | 0 | 0 |
| glt*S* | *Bacteroidetes* | 0 | 0 | 0 | 0 | 0 | 0 | 0 | 0 | 0 | 0 | 3 | 0 | 0 | 3 | 0 | 0 | 0 | 0 |
| glt*S* | *Chloroflexi* | 1 | 1 | 0 | 0 | 0 | 0 | 1 | 0 | 0 | 1 | 4 | 0 | 1 | 0 | 0 | 0 | 0 | 0 |
| glt*S* | *Cyanobacteria* | 3 | 6 | 0 | 0 | 5 | 13 | 7 | 3 | 1 | 2 | 12 | 0 | 0 | 0 | 2 | 19 | 1 | 1 |
| glt*S* | Other | 0 | 1 | 3 | 0 | 0 | 1 | 0 | 0 | 0 | 1 | 1 | 1 | 0 | 2 | 1 | 2 | 0 | 0 |
| glt*S* | Unclassified Bacteria | 0 | 4 | 0 | 1 | 0 | 4 | 0 | 1 | 2 | 8 | 5 | 1 | 0 | 6 | 0 | 11 | 5 | 2 |
| glt*S* | *Verrucomicrobia* | 0 | 0 | 0 | 0 | 0 | 0 | 2 | 0 | 0 | 0 | 3 | 0 | 0 | 1 | 0 | 0 | 0 | 0 |
| hao | *Bacteroidetes* | 0 | 0 | 0 | 0 | 0 | 0 | 0 | 0 | 0 | 0 | 1 | 0 | 0 | 0 | 0 | 0 | 0 | 0 |
| kor*AB* | *Acidobacteria* | 2 | 1 | 0 | 1 | 2 | 6 | 1 | 4 | 0 | 11 | 3 | 2 | 3 | 14 | 5 | 7 | 7 | 9 |
| kor*AB* | *Actinobacteria* | 0 | 0 | 0 | 0 | 1 | 1 | 0 | 0 | 0 | 0 | 1 | 0 | 1 | 0 | 1 | 0 | 0 | 1 |
| kor*AB* | *Bacteroidetes* | 39 | 39 | 13 | 16 | 66 | 148 | 63 | 8 | 14 | 100 | 97 | 56 | 16 | 177 | 68 | 83 | 57 | 65 |
| kor*AB* | *Chloroflexi* | 0 | 2 | 0 | 0 | 1 | 3 | 0 | 0 | 0 | 2 | 2 | 1 | 3 | 7 | 3 | 4 | 3 | 1 |
| kor*AB* | *Cyanobacteria* | 0 | 0 | 0 | 0 | 0 | 0 | 0 | 0 | 0 | 0 | 2 | 0 | 0 | 0 | 0 | 0 | 0 | 0 |
| kor*AB* | *Gemmatimonadetes* | 0 | 0 | 0 | 0 | 0 | 0 | 0 | 0 | 0 | 1 | 0 | 1 | 0 | 0 | 0 | 0 | 0 | 0 |
| kor*AB* | Other | 0 | 1 | 2 | 0 | 1 | 1 | 0 | 0 | 0 | 0 | 0 | 1 | 1 | 2 | 1 | 2 | 0 | 1 |
| kor*AB* | *Planctomycetes* | 0 | 1 | 1 | 0 | 0 | 4 | 0 | 0 | 0 | 0 | 0 | 0 | 0 | 1 | 0 | 1 | 1 | 0 |
| kor*AB* | *Proteobacteria* | 0 | 2 | 0 | 0 | 0 | 1 | 0 | 4 | 0 | 1 | 0 | 2 | 3 | 0 | 1 | 0 | 0 | 1 |
| kor*AB* | Unclassified Archaea | 6 | 0 | 0 | 0 | 1 | 4 | 2 | 0 | 0 | 2 | 0 | 1 | 1 | 1 | 5 | 0 | 0 | 1 |
| kor*AB* | Unclassified Bacteria | 10 | 13 | 0 | 9 | 19 | 25 | 12 | 9 | 3 | 32 | 39 | 15 | 7 | 57 | 17 | 18 | 37 | 24 |
| ldh | *Acidobacteria* | 0 | 1 | 0 | 0 | 0 | 0 | 0 | 0 | 0 | 0 | 0 | 0 | 0 | 0 | 0 | 0 | 0 | 0 |
| ldh | *Cyanobacteria* | 0 | 2 | 0 | 0 | 0 | 1 | 0 | 0 | 0 | 0 | 1 | 0 | 0 | 0 | 0 | 1 | 0 | 0 |
| ldh | *Gemmatimonadetes* | 0 | 0 | 0 | 0 | 0 | 0 | 0 | 0 | 0 | 0 | 0 | 0 | 0 | 1 | 0 | 0 | 0 | 0 |
| ldh | Other | 0 | 1 | 0 | 1 | 0 | 1 | 0 | 0 | 1 | 2 | 0 | 0 | 0 | 1 | 0 | 0 | 0 | 1 |
| ldh | *Planctomycetes* | 0 | 0 | 0 | 0 | 0 | 0 | 0 | 0 | 0 | 0 | 0 | 0 | 0 | 0 | 0 | 1 | 0 | 0 |
| ldh | *Proteobacteria* | 0 | 0 | 0 | 0 | 0 | 0 | 2 | 0 | 1 | 1 | 0 | 0 | 0 | 2 | 2 | 2 | 0 | 1 |
| ldh | Unclassified Bacteria | 4 | 4 | 0 | 1 | 2 | 6 | 3 | 1 | 1 | 5 | 4 | 0 | 1 | 3 | 5 | 3 | 1 | 2 |
| ldh | *Verrucomicrobia* | 0 | 0 | 0 | 0 | 0 | 0 | 0 | 0 | 0 | 1 | 0 | 0 | 0 | 0 | 0 | 0 | 0 | 0 |
| mcr*B* | Other | 0 | 0 | 0 | 0 | 0 | 0 | 0 | 0 | 0 | 0 | 0 | 0 | 0 | 1 | 0 | 0 | 0 | 0 |
| mcr*B* | Unclassified Bacteria | 0 | 0 | 0 | 0 | 0 | 0 | 0 | 0 | 0 | 1 | 0 | 0 | 0 | 0 | 0 | 0 | 0 | 0 |
| nap*A* | *Bacteroidetes* | 0 | 0 | 0 | 0 | 0 | 0 | 0 | 0 | 0 | 0 | 2 | 0 | 0 | 0 | 0 | 0 | 0 | 0 |
| nar*GH* | *Actinobacteria* | 0 | 2 | 0 | 0 | 0 | 0 | 0 | 0 | 0 | 0 | 0 | 0 | 0 | 0 | 0 | 0 | 0 | 0 |
| nar*GH* | *Proteobacteria* | 0 | 0 | 1 | 0 | 0 | 0 | 0 | 0 | 0 | 1 | 0 | 0 | 0 | 0 | 0 | 0 | 0 | 0 |
| nar*GH* | Unclassified Bacteria | 0 | 0 | 0 | 0 | 0 | 1 | 0 | 0 | 0 | 0 | 0 | 0 | 0 | 0 | 0 | 0 | 0 | 0 |
| nas | *Bacteroidetes* | 1 | 10 | 3 | 1 | 4 | 16 | 6 | 5 | 2 | 9 | 28 | 12 | 8 | 15 | 4 | 3 | 1 | 1 |
| nas | *Cyanobacteria* | 0 | 2 | 0 | 0 | 1 | 5 | 1 | 0 | 0 | 2 | 1 | 1 | 2 | 0 | 0 | 3 | 0 | 0 |
| nas | Other | 0 | 0 | 0 | 0 | 0 | 0 | 0 | 0 | 0 | 0 | 0 | 1 | 0 | 0 | 0 | 0 | 0 | 0 |
| nas | Unclassified Bacteria | 0 | 0 | 0 | 0 | 0 | 0 | 0 | 0 | 0 | 0 | 0 | 1 | 0 | 0 | 0 | 0 | 0 | 0 |
| nif*DK* | *Cyanobacteria* | 0 | 1 | 0 | 0 | 0 | 0 | 0 | 0 | 0 | 0 | 0 | 0 | 0 | 0 | 0 | 0 | 0 | 0 |
| nif*H* | *Cyanobacteria* | 1 | 0 | 0 | 0 | 0 | 4 | 0 | 0 | 0 | 0 | 0 | 0 | 0 | 0 | 0 | 0 | 0 | 0 |
| nor*BC* | *Acidobacteria* | 0 | 0 | 0 | 0 | 0 | 0 | 0 | 0 | 0 | 0 | 2 | 0 | 0 | 0 | 0 | 0 | 0 | 0 |
| nor*BC* | *Bacteroidetes* | 0 | 0 | 0 | 0 | 3 | 2 | 0 | 1 | 2 | 4 | 11 | 1 | 0 | 5 | 1 | 0 | 0 | 7 |
| nor*BC* | *Chloroflexi* | 0 | 0 | 0 | 0 | 0 | 0 | 0 | 0 | 0 | 0 | 1 | 0 | 0 | 0 | 0 | 0 | 0 | 0 |
| nor*BC* | *Cyanobacteria* | 0 | 0 | 0 | 0 | 0 | 0 | 0 | 0 | 0 | 0 | 1 | 0 | 0 | 0 | 0 | 0 | 0 | 0 |
| nor*BC* | Other | 0 | 0 | 0 | 0 | 0 | 0 | 0 | 0 | 0 | 0 | 0 | 1 | 0 | 2 | 0 | 1 | 0 | 1 |
| nor*BC* | *Proteobacteria* | 0 | 0 | 0 | 0 | 0 | 1 | 0 | 0 | 0 | 0 | 0 | 0 | 0 | 0 | 1 | 0 | 0 | 0 |
| nor*BC* | Unclassified Bacteria | 0 | 1 | 0 | 1 | 0 | 6 | 0 | 0 | 0 | 0 | 11 | 6 | 0 | 10 | 1 | 0 | 0 | 2 |
| nos*Z* | *Bacteroidetes* | 0 | 4 | 1 | 0 | 0 | 1 | 2 | 0 | 0 | 0 | 7 | 0 | 0 | 0 | 0 | 0 | 1 | 0 |
| nos*Z* | Other | 0 | 0 | 1 | 0 | 0 | 0 | 0 | 0 | 0 | 0 | 0 | 0 | 0 | 0 | 0 | 0 | 0 | 0 |
| nos*Z* | Unclassified Archaea | 0 | 0 | 0 | 0 | 0 | 0 | 0 | 0 | 0 | 0 | 0 | 0 | 0 | 0 | 1 | 0 | 0 | 0 |
| nos*Z* | Unclassified Bacteria | 0 | 1 | 0 | 0 | 0 | 0 | 0 | 0 | 0 | 1 | 3 | 0 | 0 | 0 | 0 | 2 | 0 | 0 |
| nos*Z* | *Verrucomicrobia* | 0 | 0 | 0 | 0 | 0 | 1 | 0 | 0 | 0 | 1 | 2 | 0 | 0 | 2 | 0 | 1 | 0 | 0 |
| nrf*A* | *Acidobacteria* | 0 | 0 | 1 | 0 | 0 | 0 | 0 | 0 | 0 | 0 | 1 | 0 | 0 | 0 | 0 | 0 | 0 | 0 |
| nrf*A* | *Bacteroidetes* | 0 | 0 | 0 | 0 | 0 | 0 | 0 | 0 | 0 | 0 | 3 | 0 | 0 | 0 | 0 | 0 | 0 | 0 |
| nrf*A* | Other | 0 | 0 | 2 | 0 | 0 | 0 | 0 | 0 | 0 | 0 | 0 | 0 | 0 | 0 | 0 | 0 | 0 | 0 |
| nrf*A* | *Proteobacteria* | 0 | 0 | 1 | 0 | 0 | 0 | 1 | 0 | 0 | 0 | 0 | 1 | 0 | 4 | 1 | 0 | 0 | 0 |
| nrf*A* | Unclassified Bacteria | 0 | 0 | 0 | 0 | 0 | 0 | 0 | 1 | 0 | 0 | 2 | 0 | 0 | 1 | 0 | 0 | 0 | 1 |
| prk*B* | *Actinobacteria* | 0 | 0 | 0 | 0 | 0 | 1 | 1 | 6 | 0 | 0 | 1 | 0 | 0 | 0 | 3 | 0 | 0 | 0 |
| prk*B* | *Cyanobacteria* | 0 | 2 | 0 | 0 | 4 | 8 | 2 | 1 | 0 | 2 | 1 | 1 | 0 | 0 | 0 | 3 | 0 | 0 |
| prk*B* | Other | 0 | 0 | 0 | 0 | 0 | 0 | 0 | 0 | 0 | 0 | 1 | 0 | 0 | 1 | 0 | 0 | 0 | 0 |
| prk*B* | Unclassified Bacteria | 0 | 0 | 0 | 1 | 0 | 1 | 0 | 1 | 0 | 2 | 0 | 1 | 0 | 2 | 1 | 0 | 0 | 1 |
| rbc*LS* | *Actinobacteria* | 0 | 0 | 0 | 0 | 0 | 1 | 0 | 2 | 0 | 0 | 0 | 0 | 0 | 1 | 0 | 0 | 0 | 2 |
| rbc*LS* | *Bacteroidetes* | 0 | 0 | 0 | 0 | 0 | 1 | 0 | 1 | 0 | 0 | 3 | 0 | 0 | 0 | 0 | 3 | 0 | 0 |
| rbc*LS* | *Cyanobacteria* | 0 | 2 | 0 | 0 | 1 | 6 | 2 | 0 | 2 | 0 | 2 | 1 | 0 | 0 | 0 | 4 | 0 | 0 |
| rbc*LS* | Other | 1 | 0 | 0 | 0 | 0 | 0 | 0 | 0 | 0 | 0 | 0 | 0 | 0 | 0 | 0 | 0 | 0 | 0 |
| rbc*LS* | *Planctomycetes* | 0 | 0 | 0 | 0 | 0 | 0 | 0 | 0 | 0 | 0 | 0 | 0 | 1 | 1 | 0 | 1 | 0 | 0 |
| rbc*LS* | *Proteobacteria* | 0 | 0 | 0 | 1 | 0 | 1 | 1 | 0 | 0 | 3 | 0 | 0 | 0 | 4 | 5 | 3 | 1 | 2 |
| rbc*LS* | Unclassified Archaea | 1 | 0 | 0 | 0 | 1 | 1 | 1 | 0 | 0 | 1 | 0 | 0 | 0 | 1 | 0 | 2 | 1 | 1 |
| rbc*LS* | Unclassified Bacteria | 0 | 1 | 0 | 0 | 0 | 2 | 0 | 0 | 0 | 2 | 1 | 0 | 0 | 0 | 2 | 1 | 0 | 1 |
| sse*A* | *Acidobacteria* | 0 | 1 | 0 | 0 | 0 | 0 | 0 | 1 | 0 | 0 | 1 | 0 | 2 | 4 | 0 | 0 | 0 | 0 |
| sse*A* | *Actinobacteria* | 1 | 1 | 0 | 0 | 0 | 0 | 1 | 3 | 0 | 1 | 3 | 1 | 0 | 0 | 1 | 0 | 1 | 1 |
| sse*A* | *Bacteroidetes* | 1 | 0 | 0 | 0 | 1 | 2 | 2 | 0 | 0 | 1 | 4 | 1 | 0 | 2 | 0 | 0 | 0 | 2 |
| sse*A* | *Chloroflexi* | 0 | 3 | 1 | 0 | 1 | 0 | 0 | 2 | 0 | 0 | 1 | 0 | 0 | 2 | 0 | 0 | 0 | 0 |
| sse*A* | *Cyanobacteria* | 0 | 3 | 0 | 0 | 3 | 7 | 5 | 1 | 1 | 0 | 3 | 0 | 0 | 0 | 0 | 4 | 0 | 0 |
| sse*A* | *Gemmatimonadetes* | 0 | 0 | 0 | 0 | 0 | 1 | 0 | 0 | 0 | 0 | 0 | 0 | 0 | 0 | 0 | 0 | 0 | 0 |
| sse*A* | Other | 0 | 1 | 0 | 0 | 0 | 3 | 0 | 0 | 0 | 0 | 2 | 0 | 1 | 0 | 1 | 0 | 0 | 1 |
| sse*A* | *Proteobacteria* | 0 | 0 | 0 | 0 | 0 | 1 | 0 | 0 | 0 | 0 | 0 | 0 | 0 | 1 | 0 | 0 | 0 | 0 |
| sse*A* | Unclassified Archaea | 5 | 0 | 0 | 0 | 0 | 8 | 1 | 0 | 1 | 9 | 0 | 3 | 1 | 3 | 4 | 0 | 0 | 2 |
| sse*A* | Unclassified Bacteria | 1 | 4 | 0 | 0 | 1 | 4 | 4 | 0 | 0 | 6 | 9 | 2 | 1 | 5 | 1 | 6 | 0 | 5 |
| sse*A* | *Verrucomicrobia* | 0 | 2 | 0 | 0 | 0 | 1 | 1 | 0 | 0 | 2 | 3 | 0 | 0 | 4 | 1 | 3 | 3 | 1 |

**Supplementary Table 7.** Characteristics of the 7 draft genomes reconstructed in this study.

| **Bin ID** | **Phylum/Genus** | **No. of contigs** | **Draft size (Mbp)** | **Single copy markers (out of 43)** | **Completeness (%)** | **GC content (%)** | **Contamination (%)** | **Relative Abundance (%)** |
| --- | --- | --- | --- | --- | --- | --- | --- | --- |
| Bin_62-2_MS4-1 | *Bacteroidetes/Segetibacter* | 756 | 4.9 | 36 | 92.51 | 38.4 | 1.23 | 2.1 |
| Bin_28-9_MtG-4 | *Acidobacteria/Pyrinomonas* | 119 | 3.7 | 19 | 92.25 | 45.7 | 4.27 | 5.4 |
| Bin_28-1_PT-2 | *Acidobacteria/Pyrinomonas* | 109 | 4.1 | 42 | 90.60 | 45.4 | 3.59 | 5.4 |
| Bin_33-2_MS5-1 | *Bacteroidetes/Flavisolibacter* | 76 | 3.0 | 10 | 90.18 | 41.8 | 1.23 | 0.1 |
| Bin_5-1_MS7-5 | *Acidobacteria/Pyrinomonas* | 226 | 3.2 | 29 | 84.70 | 44.3 | 4.75 | 5.4 |
| Bin_65-1_MtG22-5 | *Bacteroidetes/Niastella* | 70 | 2.0 | 9 | 62.07 | 35.7 | 0.00 | 1.1 |
| Bin_4-3_MS4-1 | *Bacteroidetes/Niastella* | 433 | 1.8 | 7 | 53.21 | 35.3 | 3.45 | 1.1 |
